# Tafazzin-Mediated Cardiolipin Remodeling Controls Metabolic Stress Response and Effector Function of Inflammatory T Cells

**DOI:** 10.1101/2025.08.25.672061

**Authors:** Xiufeng Zhao, Sophia M. Hochrein, Miriam Eckstein, Hao Wu, Katrin Sinning, Katharina J. Ermer, Sarah-Lena Puhl, Martin Eilers, Werner Schmitz, Christian Stigloher, Steven M. Claypool, Christoph Maack, Jan Dudek, Martin Vaeth

## Abstract

Clonal expansion and effector differentiation of T cells require extensive metabolic reprogramming. This includes the restructuring of the inner mitochondrial membrane (IMM) to enhance respiration by increasing chemiosmotic coupling efficiency. Cardiolipin, a unique phospholipid that is exclusively synthesized and localized in the IMM, modulates the biophysical properties of the electron transport chain (ETC) in tissues with high energy demands, such as cardiomyocytes and skeletal muscle. However, it remains unclear whether cardiolipin is also important for metabolic remodeling during T helper (Th) cell differentiation. In this study, we show that cardiolipin transacylation, catalyzed by the enzyme Tafazzin, supports the clonal expansion and effector function of inflammatory Th1 and Th17 cells *in vitro* and in models of autoimmune colitis and encephalomyelitis. At the molecular level, we demonstrate that loss of Tafazzin-mediated cardiolipin maturation induces a metabolic and transcriptional stress response in Th cells to compensate for impaired coupling efficiency of the ETC complexes and disrupted cellular redox homeostasis. However, the genetic program that restores cellular homeostasis and mitigates oxidative stress concurrently impairs the effector functions of inflammatory T cells, such as cytokine production. Our findings also provide insights into the complex clinical manifestation of patients with Barth syndrome (BTHS) caused by mutations in the human *TAFAZZIN* gene. BTHS is characterized by cardiac and skeletal myopathy as well as neutropenia and an increased susceptibility to infections. Although the molecular basis of the immunodeficiency remains poorly understood, our findings suggest that impaired Th cell function contributes to the immunopathology observed in BTHS patients.

## INTRODUCTION

Upon antigen receptor stimulation, T cells undergo extensive metabolic reprogramming to meet the elevated bioenergetic and biosynthetic demands required for cellular growth, clonal expansion and effector differentiation ^1–3^. Naïve T cells primarily rely on mitochondrial oxidative phosphorylation (OXPHOS) for efficient ATP production. In contrast, upon activation, T cells undergo a rapid metabolic shift toward aerobic glycolysis – a process reminiscent of the Warburg effect observed in cancer cells ^1,2,4^. Elevation of glycolytic flux serves fuel pentose phosphate pathway and serine metabolism required for cellular growth and proliferation. Activated T cells also exhibit enhanced mitochondrial biogenesis, remodeling of cristae structures and reorganization of the electron transport chain (ETC) to enhance OXPHOS ^4^. Beyond energy production, mitochondria serve as central hubs for pyrimidine and one-carbon metabolism, cellular signaling, stress responses and epigenetic remodeling ^4,5^. Importantly, distinct T cell subsets display specific mitochondrial profiles and metabolic dependencies – for example, memory and regulatory T cells primarily rely on OXPHOS and fatty acid oxidation, whereas effector T cells prefer to utilize aerobic glycolysis ^2,3^.

Mitochondria are highly dynamic organelles that undergo extensive remodeling during T cell activation and effector differentiation ^4^. Upon antigen receptor stimulation, transcriptional programs promote mitochondrial biogenesis, resulting in increased mitochondrial mass and enhanced respiratory capacity. Simultaneously, mitochondrial dynamics, regulated by fission and fusion, ensure proper mitochondrial distribution and function as well as the clearance of damaged organelles via mitophagy ^4–6^. Mitochondria are enclosed by a double membrane system. The outer mitochondrial membrane (OMM) forms a barrier to the cytosol, harboring proteins such as voltage-dependent anion channels (VDACs), which regulat the exchange of metabolites and proteins up to 5 kDa ^6^. In contrast, the inner mitochondrial membrane (IMM) allows the selective passage for metabolites by specific transporters and is extensively folded into cristae structures to maximize surface area. These tube-like invaginations contain the electron transport chain (ETC), metabolite transporters and regulatory protein complexes, making the IMM essential for OXPHOS, metabolic and epigenetic reprograming, signal transduction, as well as the control of various cell death pathways ^4–6^.

The IMM and OMM exhibit distinct membrane lipid compositions, reflecting their evolutionary origins ^6,7^. According to the endosymbiotic theory, the IMM is derived from the ancestral bacterial membrane, while the OMM originates from the eukaryotic membrane system ^7^. The OMM is characterized by higher levels of cholesterol and phosphatidylcholine, typical of eukaryotic membranes. In contrast, the membrane lipid cardiolipin (CL), which is found in bacteria and the IMM, accounts for ∼ 20% of the total phospholipid content in mitochondria ^8–11^. CL is unique among eukaryotic phospholipids due to its dimeric structure, consisting of four acyl chains and two phosphatidyl moieties linked by a central glycerol backbone. This distinctive architecture generates a conical shape, enabling CL to induce membrane curvatures to support cristae formation ^12^. CL directly interacts with the respiratory chain protein complexes and metabolite carriers, actively supporting the function of IMM proteins ^13–17^. Beyond its structural role, CL is essential for a range of mitochondrial functions, including mitochondrial biogenesis, fission and fusion dynamics, assembly of ETC supercomplexes (i.e., respirasomes), chemiosmotic coupling and ATP production, mitophagy, and the transport of metabolites, ions and proteins across the IMM ^10,11,18–22^. Due to its high content of unsaturated acyl chains and its proximity to the ETC – a major source of reactive oxygen species (ROS) – CL is highly susceptible to oxidative damage ^10,11^. Peroxidation of CL can disrupt its functions and, when oxidized CL species translocate to the OMM, they can initiate inflammation and apoptosis by promoting the opening of the mitochondrial permeability transition pore (mPTP) and the release of cytochrome c into the cytosol ^10,11,19^.

The biosynthesis of CL is initiated by the conversion of mitochondria or endoplasmic reticulum (ER)-derived phosphatidic acid into CDP-diacylglycerol (CDP-DAG) (reviewed in ^11,18,19^). CDP-DAG is then transported to the IMM, where it is converted into phosphatidyl glycerophosphate (PGP). PGP is subsequently dephosphorylated by protein tyrosine phosphatase mitochondrial 1 (PTPMT1) to form phosphatidylglycerol (PG). At the matrix-facing side of the IMM, nascent CL is synthesized by cardiolipin synthase 1 (CLS1) via an irreversible condensation reaction between PG and CDP-DAG. This immature CL species is characterized by highly saturated and asymmetric acyl side chains. To generate mature CL, the nascent molecule undergoes remodeling through a sequence of de-acylation and re-acylation steps. Phospholipases remove a saturated acyl chain, producing monolysocardiolipin (MLCL). Re-acylation of MLCL with unsaturated fatty acids is catalyzed by the mitochondrial transacylase Tafazzin, which is located at the outer leaflet of the IMM ^19^. Tafazzin facilitates acyl chain exchange between MLCL and other phospholipids, resulting in the production of fully remodeled, symmetric and conically shaped CL species ^10–12^. Although other enzymes, such as ALCAT1, MLCLAT and the recently discovered PLAAT1, may contribute to late-stage CL remodeling in some cell types ^10,11,19,23,24^, Tafazzin plays a unique role in mitochondrial physiology. This is evidenced by mitochondrial dysfunction in patients with Barth syndrome (BTHS), a rare X-linked recessive disorder caused by mutations in the human *TAFAZZIN* gene on chromosome Xq28 (OMIM #302060) ^25,26^.

Given the critical role of CL in mitochondrial function, patients with BTHS exhibit defects in tissues with high energetic demand, clinically manifesting as cardiomyopathy, skeletal myopathy and constitutional growth delay ^25–27^. In addition, up to 90% of BTHS patients suffer from persistent or intermittent neutropenia ^25,28^, which can be the second leading cause of death in BTHS patients ^29^. Neutropenia predisposes them to bacterial and fungal infections, including upper respiratory tract infections, chronic aphthous stomatitis, gingivitis, perianal dermatitis, and in severe cases, sepsis and multi-organ failure ^25,27^. While the exact molecular causes underlying neutropenia in BTHS remain incompletely understood ^27,28,30,31^, recent studies suggest that CL regulates neutrophil maturation and/or the clearance of senescent neutrophils ^27,30^. Although Tafazzin mutations typically do not impair basic neutrophil functions, such as phagocytosis, intracellular pathogen killing, degranulation or neutrophil extracellular trap (NET) formation, accumulating evidence indicates that elevated ER stress and unfolded protein response (UPR) signaling render Tafazzin-deficient neutrophils more susceptible to apoptosis ^27,31^.

The numbers of lymphocytes, including CD4^+^ and CD8^+^ T cell subsets, are normal in most BTHS patients ^27,32^, but whether CL also contributes to adaptive immune responses remains largely unknown. To date, only a few studies have addressed the role of CL biosynthesis and Tafazzin-mediated CL remodeling in lymphocytes ^33,34–37^. Recent studies revealed that PTPMT1-dependent *de novo* CL biosynthesis is essential for effective cytotoxic CD8⁺ T cell responses to pathogens and tumors ^34,35^. However, the contribution of Tafazzin-mediated late-stage CL remodeling remains enigmatic. Whole-body deletion of Tafazzin in mice and observations from BTHS patients revealed phenotypes that are similar to those of PTPMT1-deficient CD8⁺ T cells ^34^. However, T cell-specific ablation of Tafazzin did not recapitulate the defects observed with PTPMT1 inactivation, suggesting that this phenotype is not cell-intrinsic or is caused indirectly by alterations occurring during hematopoietic development ^27,31,34^. Moreover, whether Tafazzin and CL remodeling play a role in CD4⁺ T helper (Th) cells is currently unclear. Given that the differentiation of naïve CD4⁺ T cells into different Th cell subsets involves substantial mitochondrial biogenesis, cristae remodeling and ETC reorganization, we sought to investigate the importance of Tafazzin in inflammatory and regulatory T cells. Using two models of genetic Tafazzin inactivation – shRNA-mediated whole-body suppression of Tafazzin translation and T cell-specific ablation of the *Taz* gene – we demonstrate that CL transacylation supports metabolic stress resilience and effector function of inflammatory Th1 and Th17 cells in models of autoimmune colitis and encephalomyelitis. At the molecular level, we demonstrate that the loss of CL remodeling leads to profound morphological changes in mitochondrial cristae morphology, along with metabolic and transcriptional stress responses to cope with mitochondrial insufficiency and disrupted cellular redox homeostasis. These findings reveal a critical role for Tafazzin-mediated CL remodeling in supporting the effector function of Th cells, potentially contributing to the immunodeficiency and infection susceptibility observed in patients with BTHS.

## RESULTS

### Cardiolipin composition changes dynamically during CD4^+^ T cell differentiation

The differentiation of naïve T cells into inflammatory and regulatory subsets involves significant mitochondrial remodeling, respirasome reorganization at the IMM and an overall increase in OXPHOS. Although nearly all mitochondrial membrane-associated processes depend on CL, the impact of CL alterations in mitochondrial remodeling during T cell differentiation remains largely unknown. To investigate whether T cell activation and effector differentiation affects the amount and composition of CL in different CD4^+^ T cell subsets, we polarized naïve *wild type* (WT) T cells into inflammatory T helper 1 (Th1) and Th17 cells, as well as T regulatory (Treg) cells (Fig. S1A).

To determine the mitochondrial lipid composition in naïve and differentiated CD4^+^ T cells, we employed targeted reverse-phase liquid chromatography coupled with mass spectrometry (LC/MS) and analyzed 10^7^ T cells of each subset (Fig. 1 and Fig. S1B-D). Intriguingly, the relative amounts of both CL and monolysocardiolipin (MLCL) significantly decreased in differentiated Th1, Th17 and induced T regulatory (iTreg) cells compared to naïve T cells (Fig. 1A and Fig. S1B). However, this decrease may be attributed to the overall accumulation of other lipid species during T cell activation. Notably, the CL-to-MLCL ratio increased in differentiated Th cells, suggesting that activation and differentiation enhances CL maturation in T cells (Fig. 1B). The predominant CL species in naïve T cells, CL(72:08), CL(74:08), CL(74:09) and CL(74:10), accounting for ∼ 70% of their total mitochondrial CL content, were markedly downregulated in Th1, Th17 and iTreg cells (Fig. S1C). In contrast, CL(68:04), CL(68:05), CL(68:06) and CL(70:05), which were nearly undetectable in naïve T cells, significantly increased in differentiated T cells (Fig. 1C and Fig. S1D). These findings demonstrate that the abundance and composition of mitochondrial lipid species undergo dynamic changes during effector T cell differentiation (Fig. 1D-F). Furthermore, Th1, Th17 and iTreg cells exhibit distinct CL species compositions, suggesting that late-stage CL remodeling plays subset-specific roles in inflammatory and regulatory T cell subsets.

**Figure 1.**
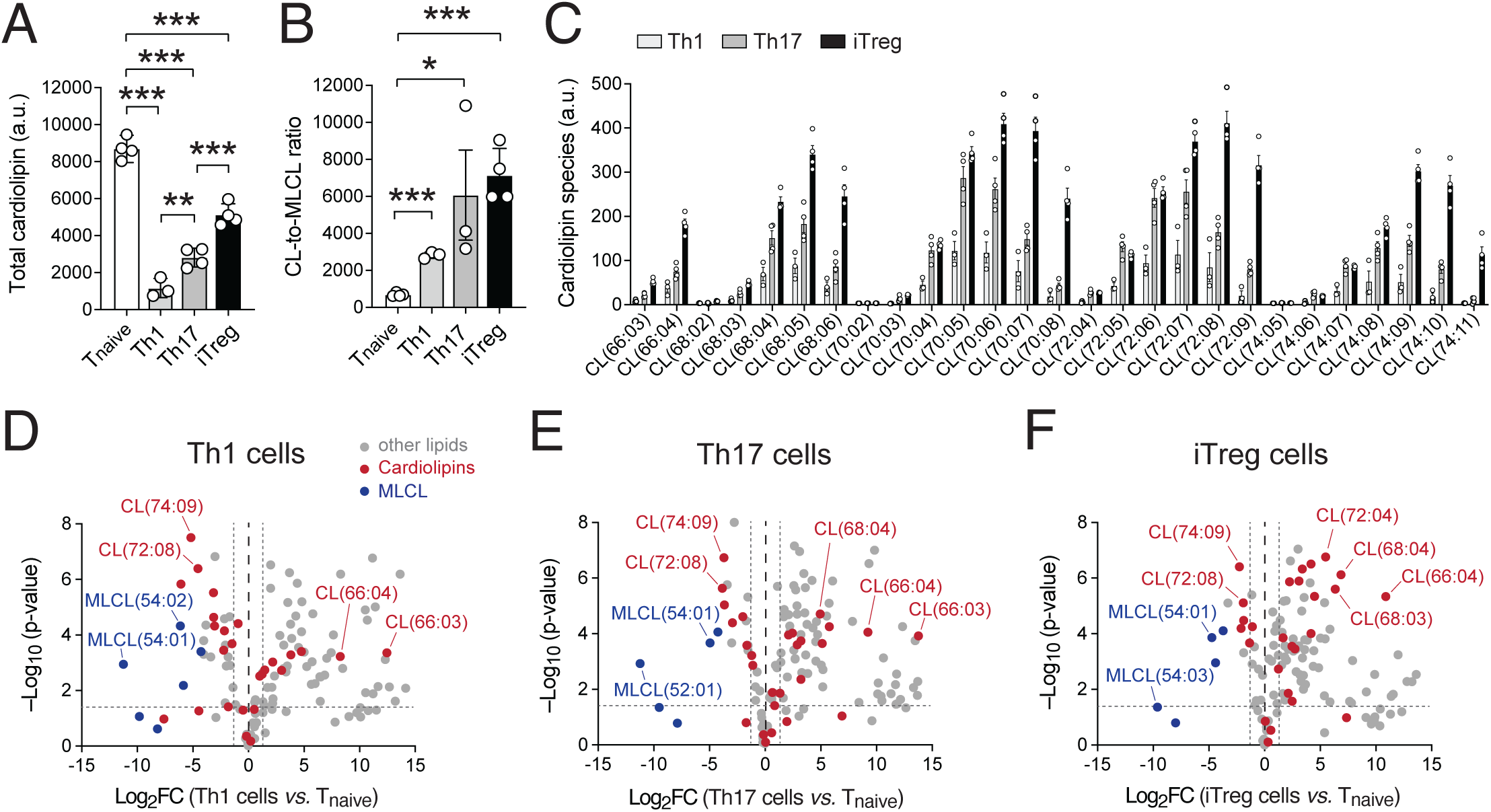
Dynamic changes in cardiolipin composition during CD4^+^ T cell differentiation. **(A)** Analysis of total cardiolipin (CL) content per 1×10^6^ cells in naïve CD4^+^ T cells and after *in vitro* differentiation into Th1, Th17 and iTreg cells by liquid chromatography and mass spectrometry (LC/MS); 3-4 biological replicates per group. **(B)** Ratio of mature CL to monolysocardiolipin (MLCL) levels in naïve, Th1, Th17 and iTerg cells; 4 biological replicates per group. **(C)** Analysis of individual cardiolipin species in differentiated Th1, Th17 and iTreg cells using LC/MS; numbers in brackets indicate number of carbon atoms to unsaturated C=C bonds of the acyl side chains. **(D-F)** Volcano plot analyses of differential lipid species in Th1 (C), Th17 (D) and iTreg (E) cells compared to naive CD4^+^ T cells; 4 biological replicates per group. CL and MLCL lipid species are depicted in red and blue, respectively. Statistical analyses in (A) and (B) by unpaired Student’s t-tests. *, p<0.05; **, p<0.01, ***, p<0.001.

### Suppression of *Tafazzin* impairs the effector function of inflammatory T cells

Having established that the amount and composition of CL is modulated during Th cell differentiation (Fig. 1 and Fig. S1), we next sought to investigate the physiological significance of CL remodeling in inflammatory and regulatory T cells. A well-established model to study late-stage CL remodeling through Tafazzin-mediated transacylation *in vivo* is a transgenic mouse line expressing a doxycycline-inducible short hairpin (sh) RNA cassette, enabling systemic *Tafazzin* mRNA knockdown ^38,39^. After doxycycline treatment for 6-8 months, *Rosa26^H1/tetO-shRNA:TAZ^* mice (hereafter referred to as *Taz*^KD^ mice) resemble the clinical hallmarks of BTHS, including cardiomyopathy and skeletal myopathy, along with characteristic physiological, biochemical and ultrastructural abnormalities observed in BTHS patients^38–40^.

We first assessed *Tafazzin* gene expression in naïve and anti-CD3/CD28-stimulated CD4^+^ T cells isolated from *Taz*^KD^ mice that had been maintained on a doxycycline diet for three months. Both naïve and activated T cells showed a robust ∼ 60-80% reduction in *Tafazzin* expression compared to WT controls (Fig. 2A), making these mice a suitable model to study the role of Tafazzin-mediated CL remodeling during T cell differentiation. Three months old *Taz*^KD^ mice had normal numbers of immature, CD4^+^CD8^+^ double positive (DP), CD4^+^ and CD8^+^ single positive (SP) thymic lymphocytes after doxycycline treatment (Fig. S2A,B). Furthermore, *Tafazzin* depletion did not alter the composition of naïve, effector and/or memory CD4^+^ and CD8^+^ T cell subsets (Fig. S2C-E) or affect the number of Foxp3^+^ Treg cells in peripheral lymphoid organs (Fig. S2F). Of note, *Taz*^KD^ mice fed with doxycycline for more than eight months developed a fulminant BTHS pathology, which was accompanied with reduced Treg frequencies and shifts in their effector/memory T cells (data not shown). Thus, long-term systemic suppression of *Tafazzin* may have indirect effects that could confound T cell composition and/or function in diseased *Taz*^KD^ mice. To minimize the secondary effects of extended and systemic *Tafazzin* inhibition, we isolated CD4^+^ T cells from *Taz*^KD^ mice that were maintained on doxycycline diet for three months and analyzed T cell differentiation and effector function *in vitro* (Fig. 2B-H and Fig. S2G-I). Knockdown of *Tafazzin* did not affect the upregulation of activation markers (Fig. 2B and Fig. S2G), proliferation (Fig. S2H) or the viability of Th1, Th17 and iTreg cells *in vitro* (Fig. S2I). However, *Taz*^KD^ T cells showed a significantly decreased expression of the Th1 and Th17 cell ‘signature’ cytokines IFNγ and IL-17A/F, respectively (Fig. 2C-F). In contrast, Foxp3 expression and iTreg cell differentiation remained unaffected, suggesting that Tafazzin-mediated CL remodeling supports the inflammatory, but not the regulatory, functions of CD4^+^ T cells.

**Figure 2.**
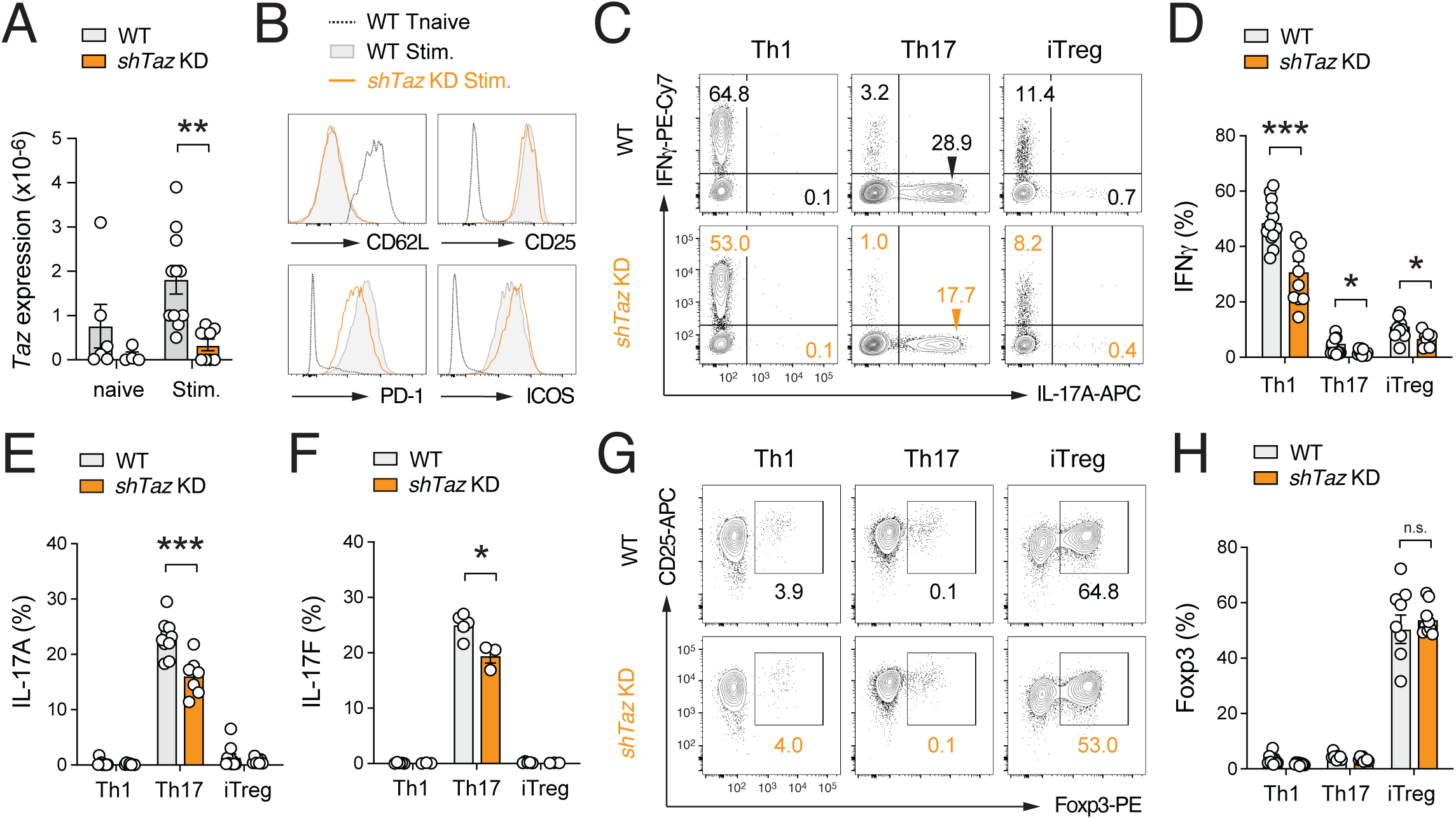
Cardiolipin remodeling supports the effector function of inflammatory T cells. **(A)** Analysis of *Tafazzin* (*Taz*) gene expression in naïve and anti-CD3/CD28 stimulated CD4^+^ T cells of WT and *Taz*^KD^ mice by qRT-PCR; means ± SEM of 4-11 mice. **(B)** Representative flow cytometric analysis of CD62L, CD25, PD-1 and ICOS protein expression on naïve and anti-CD3/CD28 activated T cells of WT and *Taz*^KD^ mice. **(C-F)** Flow cytometric analysis of IFNγ **(D),** IL-17A **(E)** and IL-17F **(F)** expression of Th1, Th17 and iTreg cells after re-stimulation with PMA/Iono for 5 h. CD4^+^ T cells were differentiated *in vitro* from naïve T cells of WT and *Taz*^KD^ mice; means ± SEM of 3-13 mice. **(G** and **H)** Foxp3 expression in WT and *Taz*^KD^ T cells differentiated under Th1, Th17 and iTreg-polarizing conditions; means ± SEM of 9 mice. Statistical analyses in (A), (D), (E), (F) and (H) by unpaired Student’s t-tests. *, p<0.05; **, p<0.01, ***, p<0.001.

### T cell-specific ablation of Tafazzin impairs Th1 and Th17 cell responses

Th1 and Th17 cells differentiated *in vitro* from naïve T cells of doxycycline-treated *Taz*^KD^ mice produced significantly less IFNγ, IL-17A and IL17F (Fig. 2C-F). However, the efficiency of shRNA-mediated *Tafazzin* suppression was incomplete (∼ 60-80% reduction) (Fig. 2A) and the systemic knockdown of the *Tafazzin* gene in *Taz*^KD^ mice may have indirect effects on T cells. Furthermore, doxycycline administration may induce microbial dysbiosis in mice ^41^, which could lead to both direct and indirect effects on the immune system ^42^.

To eliminate potential confounding effects of doxycycline, we generated mice with T cell-specific, genetic deletion of the *Tafazzin* gene by crossing *Taz*^fl/fl^ mice ^43^ with *Cd4*^Cre^ animals. In *Taz*^fl/fl^*Cd4*^Cre^ mice, *Tafazzin* is ablated at the late CD4^+^CD8^+^ DP stage of thymic T cell development, enabling the study of Tafazzin-mediated CL remodeling specifically in peripheral T cells without any indirect effects associated with systemic BTHS pathology. As expected, *Tafazzin* gene (Fig. 3A) and protein (Fig. 3B) expression were completely abolished in T cell subsets of *Taz*^fl/fl^*Cd4*^Cre^ mice. *Taz*^fl/fl^*Cd4*^Cre^ mice were born at expected Mendelian ratios and had normal numbers of immature, DP and SP thymic T cells compared to littermate WT controls (Fig. S3A,B). The frequency and number of both conventional and regulatory T cell subsets were unchanged and the *Taz*^fl/fl^*Cd4*^Cre^ mice showed no obvious immune dysregulation, such as an accumulation of effector/memory T cells in their peripheral lymphoid organs (Fig. S3C-F). Importantly, mature CL was virtually absent in Th1 and Th17 cells, while MLCL levels were markedly increased (Fig. 3C). This was not caused by a decrease in specific CL species, but rather by the global absence of CL in Tafazzin-deficient Th1 and Th17 cells (Fig. S3G,H). These data demonstrate that genetic ablation of Tafazzin effectively abolishes CL transacylation in T cells. As observed in T cells from *Taz*^KD^ mice, complete loss of Tafazzin did not affect the expression of activation markers (Fig. S3I) or the viability of Th1, Th17 and iTreg cells *in vitro* (Fig. S3J), but delayed cell cycle entry (Fig. 3D) and proliferation (Fig. 3E). The expression of the lineage-defining transcription factors T-bet (Fig. S3K) and RORγt (Fig. S3L) were comparable between WT and Tafazzin-deficient Th1 and Th17 cells, respectively, indicating that the differentiation of these subsets was not impaired. In contrast, the production of IFNγ and IL-17A was significantly reduced in Tafazzin-deficient T cells compared to WT controls (Fig. 3F,G). Foxp3 expression remained unaltered (Fig. 3H), suggesting that the differentiation of regulatory T cells was not affected. Consistent with the findings in *Taz*^KD^ mice, T cell-specific ablation of Tafazzin further supports the notion that CL remodeling is critical for the inflammatory function of Th1 and Th17 cells.

**Figure 3.**
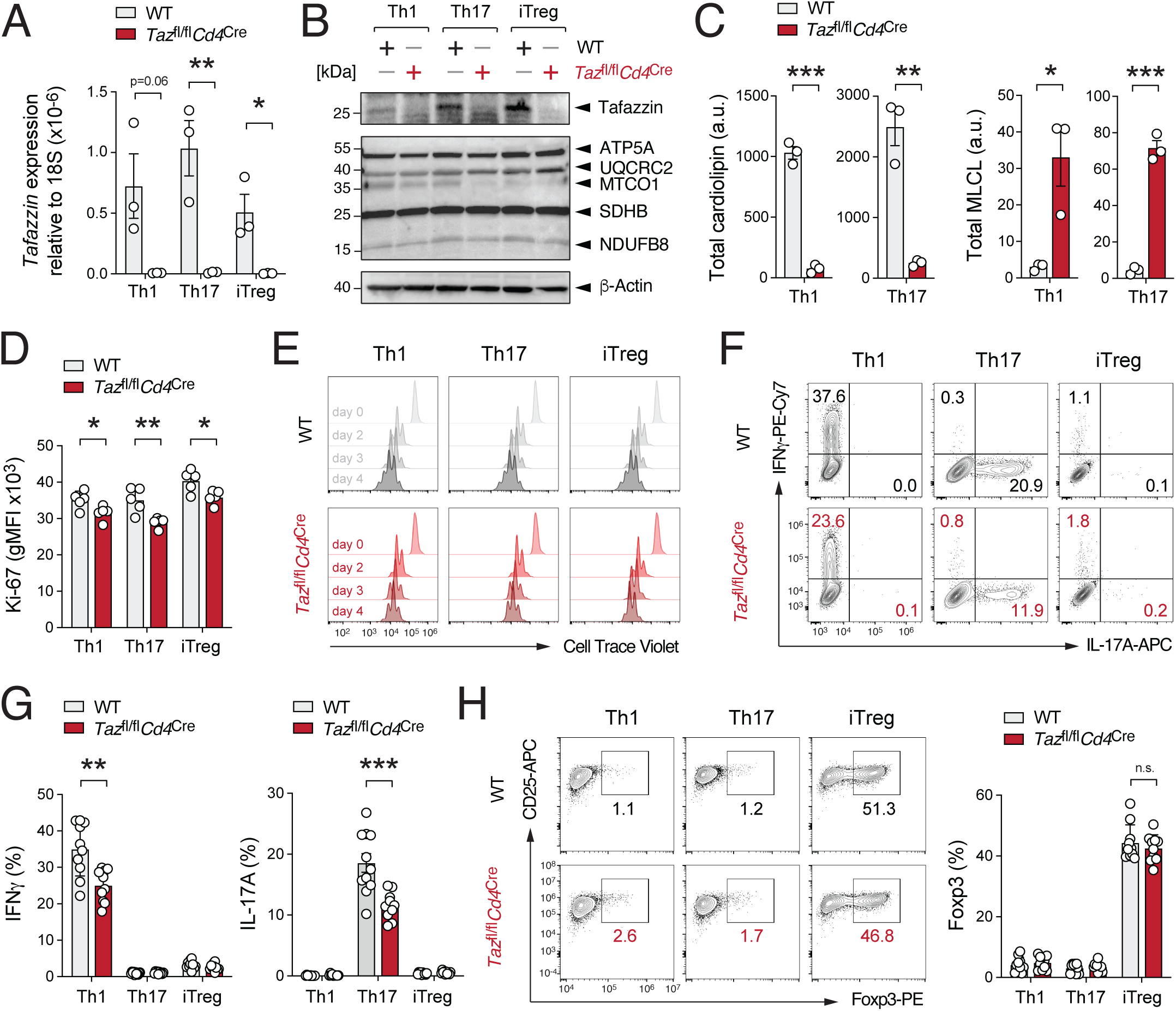
T cell-specific ablation of Tafazzin limits Th1 and Th17 cell responses. **(A)** Analysis of *Tafazzin* gene expression in differentiated Th1, Th17 and iTreg cells of WT and *Taz*^fl/fl^*Cd4*^Cre^ mice by qRT-PCR; means ± SEM of 3 mice. **(B)** Immunoblot analysis of Tafazzin and electron transport chain (ETC) protein expression in Th1, Th17 and iTreg cells of WT and *Taz*^fl/fl^*Cd4*^Cre^ mice. **(C)** Analysis of total cardiolipin (CL) and monolysocardiolipin (MLCL) content per 1×10^6^ Th1 and Th17 cells using liquid chromatography coupled with mass spectrometry (LC/MS) analysis; means ± SEM of 3 mice. **(D)** Flow cytometric analysis of Ki-67 expression in WT and Tafazzin-deficient T cells cultured for 3 days under Th1, Th17 and iTreg culture conditions; means ± SEM of 5 mice. **(E)** Proliferation analysis of WT and Tafazzin-deficient Th1, Th17 and iTreg cells by cell trace violet (CTV) dilution over a course of 4 days after anti-CD3/CD28 stimulation. **(F** and **G)** Flow cytometric analysis of IFNγ and IL-17A production by WT and Tafazzin-deficient Th1, Th17 and iTreg cells; means ± SEM of 10-11 mice. **(H)** Foxp3 expression in WT and Tafazzin-deficient T cells differentiated under Th1, Th17 and iTreg-polarizing conditions; means ± SEM of 10-11 mice. Statistical analyses in (A), (C), (D), (G) and (H) by unpaired Student’s t-tests. *, p<0.05; **, p<0.01, ***, p<0.001.

### Ablation of Tafazzin in T cells ameliorates autoimmunity *in vivo*

Both Th1 and Th17 cells promote immunopathology in various autoimmune diseases, including multiple sclerosis (MS) and inflammatory bowel disease (IBD). To investigate the role of Tafazzin in Th1 and Th17 cell *in vivo*, we utilized two animal models of T cell-mediated inflammation; experimental autoimmune encephalomyelitis (EAE) (Fig. 4A-G) and adoptive transfer colitis (Fig. 4H-K), which resemble clinical characteristics of human MS and IBD, respectively.

**Figure 4.**
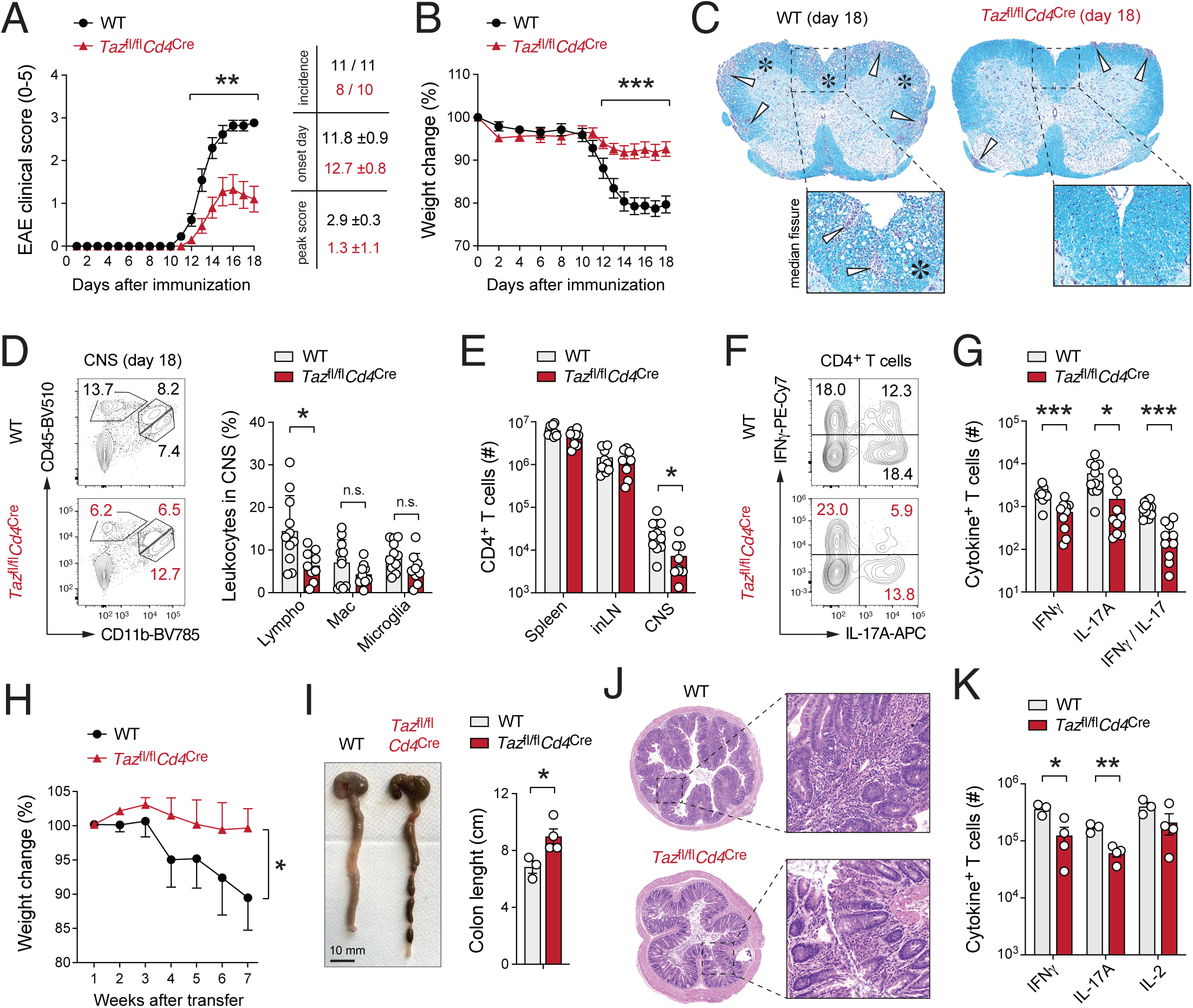
Ablation of Tafazzin alleviates T cell-mediated autoimmunity. **(A**-**D)** Myelin oligodendrocyte (MOG)-induced experimental autoimmune encephalomyelitis (EAE) is ameliorated in *Taz*^fl/fl^*Cd4*^Cre^ mice. **(A** and **B)** Clinical EAE scores (A) and relative weight changes (B) of WT and *Taz*^fl/fl^*Cd4*^Cre^ mice after immunization with MOG_35-55_ peptide emulsified in CFA; means ± SEM of 10-11 mice per cohort. **(C)** Representative histopathological examination of caudal spinal cord sections of WT and *Taz*^fl/fl^*Cd4*^Cre^ mice 18 days after MOG_35-55_ peptide immunization. White arrows and asterisks indicate leukocytic infiltrates and areas of demyelination, respectively. **(D)** Flow cytometric analysis of immune cell populations in the CNS of WT and *Taz*^fl/fl^*Cd4*^Cre^ mice after immunization with MOG_35-55_ peptide; means ± SEM of 10 mice per cohort. **(E)** Absolute CD4^+^ T cell numbers in the spleen, inguinal lymph nodes (inLN) and CNS of WT and *Taz*^fl/fl^*Cd4*^Cre^ mice after immunization with MOG_35-55_ peptide; means ± SEM of 10 mice per group. **(F** and **G)** Frequencies (F) and absolute cell numbers (G) of IFNγ and IL-17A-producing CD4^+^ T cells in the CNS of WT and *Taz*^fl/fl^*Cd4*^Cre^ mice 18 days after MOG_35-55_ immunization; means ± SEM of 10 mice. **(H**-**K)** Tafazzin-deficient T cells fail to induce autoimmune colitis after transfer into lymphopenic recipient mice. **(H)** Weight loss of *Rag1*^-/-^ host mice after transfer of naive CD4^+^ T cells from WT or *Taz*^fl/fl^*Cd4*^Cre^ mice; means ± SEM of 6-8 host mice. **(I)** Representative macroscopic pictures (left) and colon lengths (right) of *Rag1*^-/-^ mice 9 weeks after T cell transfer; means ± SEM of 3-4 recipient mice. **(J)** Representative H&E-stained colon sections detecting inflammatory tissue damage and leukocyte infiltration. **(K)** Absolute numbers of IFNγ and IL-17A-producing CD4^+^ T cells in the intestine of *Rag1*^-/-^ recipient mice 7 to 9 weeks after transfer of WT or Tafazzin-deficient T cells; means ± SEM of 3-4 host mice. Statistical analyses in (D), (E), (G) and (K) by unpaired Student’s t-tests. Statistical analyses shown in (A, B) by Mann-Whitney-U-tests. *, p<0.05; **, p<0.01, ***, p<0.001.

We induced EAE in *Taz*^fl/fl^*Cd4*^Cre^ and littermate control mice by immunization with MOG_35-55_ peptide and monitored disease progression over 18 days (Fig. 4A,B). MOG-reactive Th1 and Th17 cells initiate an autoimmune response that mimics MS by inducing central nervous system (CNS) inflammation, recruiting macrophage and neutrophil and promoting neuronal demyelination. Mice with T cell-specific deletion of Tafazzin were largely protected from EAE-induced immunopathology, as evidenced by reduced paralysis of the extremities (Fig 4A), less inflammation-induced weight loss (Fig. 4B) and diminished demyelination of the spinal cord (Fig. 4C) as clinical readouts for the EAE immunopathology. The number of CNS-resident lymphocytes was significantly reduced in *Taz*^fl/fl^*Cd4*^Cre^ mice compared to WT control animals (Fig. 4D and S4A), primarily due to lower infiltration of CD4^+^ T cells (Fig. 4E). Although the expression levels of IFNγ, IL-17A and GM-CSF in CNS-infiltrating T cells were only slightly altered in *Taz*^fl/fl^*Cd4*^Cre^ mice (Fig. S4B), the numbers of IFNγ and IL-17A-producing encephalitogenic CD4^+^ T cells were significantly reduced (Fig. 4G).

In a second model, we investigated the role of Tafazzin in Th1 and Th17 cells by transferring naïve CD45RB^hi^ T cells from WT and *Taz*^fl/fl^*Cd4*^Cre^ mice into lymphopenic (*Rag1*^-/-^) host mice to induce autoimmune colitis (Fig. 4H-K). In the absence of Foxp3^+^ Treg cells, transplanted naïve CD4^+^ T cells differentiate into colitogenic Th1 and Th17 cells, which provoke a chronic inflammation in both the small and large intestine. Disease progression was monitored by assessing weight loss and diarrhea over the course of 7-9 weeks (Fig. 4H-J). Mice receiving Tafazzin-deficient T cells exhibited reduced weight loss compared to animals that were transplanted with WT lymphocytes (Fig. 4H). Macroscopic inspection of colonic samples of mice that were transplanted with WT T cells revealed pronounced colonic shortening, thickening and edema (Fig. 4I). However, mice that received Tafazzin-deficient T cells showed no macroscopic signs of colitis (Fig. 4I). Histopathologic examination of colonic tissue sections revealed significantly milder IBD symptoms in mice after transfer of Tafazzin-deficient T cells, characterized by moderate lymphocytic infiltration and limited goblet cell depletion (Fig. 4J). In contrast, transfer of WT T cells in *Rag1*^-/-^ mice caused severe colonic inflammation, marked by massive lymphocytic infiltration, epithelial hyperplasia, goblet cell depletion and ulceration (Fig. 4J). The numbers of Tafazzin-deficient CD4^+^ T cells in the small intestine and colon were significantly lower than those in WT controls (Fig. S4C), while Foxp3^+^ Treg cell numbers were comparable (Fig. S4D). Consistent with reduced colitis immunopathology, *Rag1*^-/-^ mice transplanted with Tafazzin-deficient T cells had fewer cytokine-producing T cells (Fig. 4K) and neutrophils (Fig. S4E) in their intestines compared to WT controls.

Together, these data demonstrate that Tafazzin-mediated CL remodeling is critical for the clonal expansion and effector function of encephalitogenic and colitogenic Th1 and Th17 cells in models of neuroinflammation and autoimmune colitis.

### Tafazzin controls metabolic stress response pathways in T cells

To gain a deeper understanding of how Tafazzin-mediated CL remodeling affects inflammatory T cells at the molecular level, we conducted transcriptomic (Fig. 5A-E and Fig. S5A-G) and metabolomic analyzes (Fig. 5H-J) using RNA sequencing (RNA-seq) and liquid chromatography with mass spectrometry (LC/MS), respectively. The expression of the *Tafazzin* gene was almost undetectable in all three Tafazzin-deficient T cell subsets (Fig. S5A-D). The transcriptome profiles of WT and Tafazzin-deficient T cell subsets were markedly altered, revealing 261, 119 and 241 differentially expressed genes (DEGs) in Th1 (Fig. S5A), Th17 (Fig. S5B) and iTreg cells (Fig S5C), that were significantly (padj < 0.01) up-or downregulated. Among them, 42 DEGs were shared by all three T cell subsets (Fig. 5A), suggesting that Tafazzin controls both a common transcriptional signature and genes that contribute to subset-specific programs of Th1, Th17 and iTreg cells. The common DEGs also included *Tafazzin*, which was strongly downregulated in all Tafazzin-deficient T cell subsets, as expected (Fig. S5D). Among the DEGs were also numerous metabolic genes, including amino acid transporters (e.g., *Slc1a4, Slc1a5*, *Slc6a9, Slc7a3, Slc7a11, Slc38a3*), components of the respiratory chain (*Cox6a, Cyb5r1, Cyb5r2*) and enzymes involved in lipid biosynthesis and fatty acid metabolism (*Fads2, Acsl6, Elovl6, Acot2, Paqr3, Plppr3, Pcyt2*) (Fig. 5A and Fig. S5A-C). Of note, the lineage-defining transcription factors *Tbx21* (encoding T-bet), *Rorc* (RORγt) and *Foxp3* were expressed at similar levels in WT and Tafazzin-deficient Th1, Th17 and iTreg cells (Fig. S5E).

**Figure 5.**
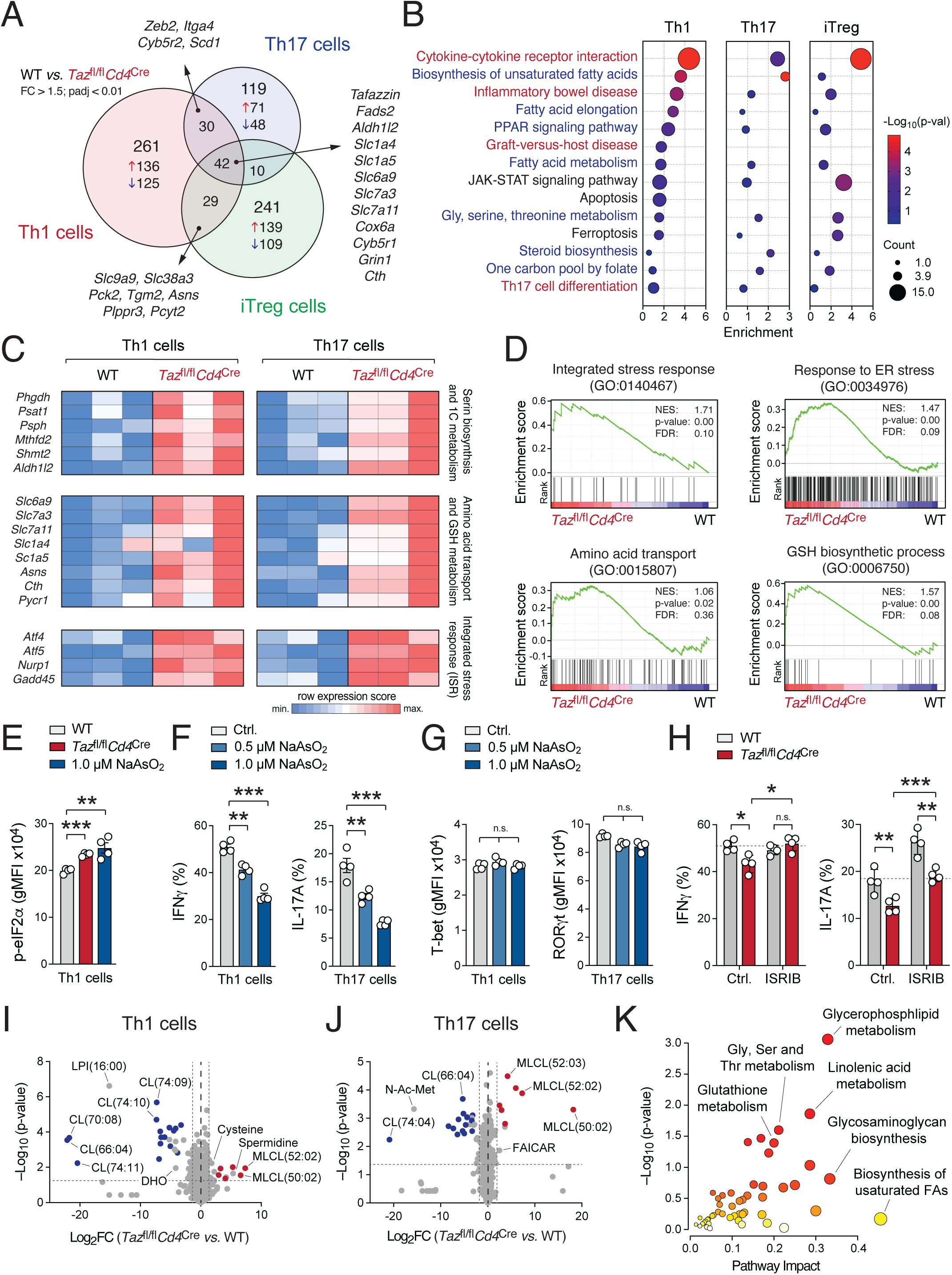
Transcriptional and metabolic consequences of Tafazzin ablation in T cells. **(A)** Venn diagram of > 1.5-fold DEG (padj < 0.01) in Tafazzin-deficient Th1, Th17 and iTreg cells. Numbers of up-and downregulated genes are displayed by red and blue arrows, respectively. **(B)** Kyoto Encyclopedia of Genes and Genomes (KEGG) pathway enrichment analysis of total DEG between WT and Tafazzin-deficient Th1, Th17 and iTreg cells; metabolic and immunological gene signatures are depicted in blue and red, respectively. **(C)** Heatmap analysis of selected genes involved in *serine and one-carbon metabolism*, *amino acid transport*, *glutathione biosynthesis* and *integrated stress response* in WT and Tafazzin-deficient Th1 and Th17 cells. **(D)** Gene set enrichment analyses (GSEA) of *glutathione biosynthesis* process, *integrated stress response*, *response to ER stress* and *amino acid transport* gene signatures. **(E)** Analysis of eIF2α phosphorylation in WT and Tafazzin-deficient T cells and WT T cells treated with 1 µM NaAsO_2_ by flow cytometry; means ± SEM of 4 mice. **(F)** Flow cytometric analysis of IFNγ and IL-17A expression after in Th1 and Th17 cells treated with or without NaAsO_2_ for 48h; means ± SEM of 4 mice. **(G)** Flow cytometric analysis of T-bet and RORγt protein expression in Th1 and Th17 cells treated with or without NaAsO_2_ for 48h; means ± SEM of 4 mice. **(H)** Analysis of IFNγ and IL-17A expression in WT and Tafazzin-deficient Th1 and Th17 cells treated with and without 500 nM ISRIB; means ± SEM of 4 mice. **(I-K)** Metabolomic profiling of WT and Tafazzin-deficient Th1 and Th17 cells. **(I** and **J)** Volcano plots of differential metabolite levels between WT *versus* Tafazzin-deficient Th1 (I) and Th17 cells (J) analyzed by liquid chromatography and mass spectrometry (LC/MS); 3 biological samples per group. Cardiolipin (CL) and monolysocardiolipin (MLCL) species are labelled in red and blue, respectively. **(K)** Metabolic network analysis using differential metabolite concentrations (p < 0.05) and DEGs (p < 0.01) between WT and Tafazzin-deficient T cells.

To better understand how CL remodeling regulates biological processes in Th1, Th17 and iTreg cells, we performed unbiased pathway enrichment analyses using the Kyoto Encyclopedia of Genes and Genomes (KEGG) database and gene set enrichment analyses (GSEA) (Fig. 5B-D). These analyses revealed that CL plays a role not only in immunological pathways such as *cytokine receptor interaction*, *inflammatory bowel disease* and *Th17 cell differentiation* but also in lipogenic metabolic pathways, including *biosynthesis of unsaturated fatty acids*, *fatty acid metabolism* and *steroid biosynthesis* (Fig. 5B). A detailed examination of the DEGs revealed that deletion of Tafazzin leads to the upregulation of genes involved in one-carbon metabolism, integrated stress response (ISR) and response to ER stress (Fig 5C,D). Additionally, genes involved in amino acid metabolism and glutathione biosynthesis were significantly increased (Fig. 5C,D). The ISR is a conserved cellular signaling pathway that enables cells to adapt to various (metabolic) stressors, including ER stress (Fig. 5D), amino acid deprivation (Fig. 5D) and oxidative stress (Fig. S5F). Activation of the ISR is initiated by phosphorylation of eukaryotic initiation factor 2 (eIF2α), leading to the induction of activating transcription factor (ATF) family members ^44^. Consequently, we observed increased phosphorylation of eIF2α in Tafazzin-deficient T cells compared to WT controls, to a similar extent as seen in WT T cells treated with the ISR-inducing agent sodium arsenite (NaAsO_2_) (Fig. 5E). We also found enhanced expression of the ISR-inducible transcription factors ATF4 and ATF5 (Fig 5C), along with a significant enrichment in the *ATF4 activated genes in response to ER stress* gene signature (Fig. S5G). Notably, approximately 20% of the genes regulated by Tafazzin in CD4^+^ T cells appear to be direct targets of ATF4, based on the overlap of DEGs in WT *versus* Tafazzin-deficient T cells and genome-wide ATF4 binding sites identified by chromatin immunoprecipitation with DNA sequencing (ChIP-seq) analyses in T cells ^45^ (Fig. S5H). Although we observed significantly reduced protein expression of IFNγ, IL-17A and IL-17F (Fig. 3F,G), transcription of their corresponding genes remained unchanged (Fig. S5I). A recent study by Asada *et al.* revealed that the ISR regulates cytokine production by modulating cytokine mRNA translation ^46^. Consistently, treatment of WT Th1 and Th17 cells with NaAsO_2_ mimicked the phenotype of Tafazzin-deficient T cells, resulting in impaired IFNγ and IL-17A protein expression (Fig. 5F) without affecting their differentiation (Fig. 5G). Importantly, treatment of Tafazzin-deficient Th1 and Th17 cells with ISRIB – a small molecule inhibitor of eIF2α activity – restored IFNγ and IL-17A expression to WT levels (Fig. 5H). Moreover, induction of ISR in WT cells using NaAsO_2_ led to reduced Ki-67 expression (Fig. S5J), while ISRIB increased its expression in Tafazzin-deficient T cells (Fig. S5K). Collectively, these findings revealed that CL deficiency activates the ISR pathway, resulting in ATF4-dependent gene expression and metabolic reprogramming, while suppressing cap-dependent cytokine mRNA translation.

To further investigate how changes in gene expression and translation impact cellular metabolism in Tafazzin-deficient Th1 and Th17 cells, we performed targeted metabolomic analyses of both polar and lipid fractions using LC/MS (Fig. 5I,J and Fig. S5I,J). As expected, the metabolic signatures of Th1 (Fig. 5I) and Th17 cells (Fig. 5J) were predominated by the depletion of CL and an enrichment of different MLCL species. In addition, various metabolic intermediates involved in the biosynthesis of pyrimidines and purines (including *carbamoyl-aspartate dihydroorotate* [DHO], *orotate*, *5-formamidoimidazole-4-carboxamide ribotide* [FAICAR]) and the metabolism of polyamines (*spermidine, acetyl-putrescine*), amino acids (*cysteine, kynurenate, N-acetyl-aspartate* [NAA]) and lipids (*acylcarnitine* [AC], *phosphatidylethanolamine* [PEA], *phosphatidylcholine* [PC], *bis(monoacylglycerol)phosphate* [BMP/PG], *phosphatidylinositol* [PI] and *lysophoshatidylcholine* [LPC]) were significantly dysregulated in Tafazzin-deficient Th1 and Th17 cells (Fig. 5I,J and Fig. S5L,M). To further elucidate the metabolic impact of *Tafazzin* deletion in T cells, we performed metabolite set enrichment analysis (MSEA) (Fig. S5K) and conducted joint pathway correlation analysis by integrating our metabolomic and transcriptomic datasets (Fig. 5K). These analyses confirmed that Tafazzin ablation not only disrupts the biosynthesis of CL glycerophospholipids but also exerts broad metabolic effects, impacting amino acid, glutathione and lipid metabolism in inflammatory CD4^+^ T cells (Fig. 5K).

### Cardiolipin remodeling controls mitochondrial structure and ROS production in T cells

Tafazzin-mediated CL transacylation regulates mitochondrial biogenesis and dynamics, as well as transport across the IMM and mitochondrial respiration in cardiac myocytes and other (non-lymphoid) tissues ^8,9^. However, whether CL remodeling plays a similar role in CD4^+^ T cells remains unclear. To address this question, we investigated the impact of Tafazzin ablation on mitochondrial function in Th1, Th17 and iTreg cells using a Seahorse extracellular flux analyzer (Fig. S6A-C). Surprisingly, however, we did not find an impairment of the basal or the maximal oxygen consumption rate (OCR), after uncoupling with the protonophore FCCP (Fig. 6SA,B). We did also not observe changes in the spare respiratory capacity (SRC) in Tafazzin-deficient T cells compared to WT controls (Fig. 6SB). Furthermore, ATP-linked respiration – measured following inhibition of complex V (ATP synthase) by oligomycin – remained unchanged in Tafazzin-deficient Th1, Th17 and iTreg cells (Fig. S6C). Total cellular ATP levels were also unaffected in absence of mature CL (Fig. S6D), suggesting that Tafazzin-deficient T cells do not experience a global bioenergetic deficit. However, the coupling efficiency (Fig. 6A) – defined as the proportion of oxygen consumption by ATP-linked respiration *versus* the proton leak across the IMM (Fig. S6E) – was significantly reduced in Tafazzin-deficient T cells, indicating that chemiosmotic coupling of ETC complexes is impaired in the absence of CL. To directly assess substrate utilization at individual ETC complexes, we employed an alternative extracellular flux protocol using permeabilized Th1 and Th17 cells, along with complex-specific substrates and inhibitors. When pyruvate was provided as a substrate followed by inhibition of complex I with rotenone, OCR at complex I was elevated in Tafazzin-deficient Th1 and Th17 cells (Fig. 6B,C). In contrast, respiration through complexes II and III was significantly reduced in Tafazzin-deficient T cells, as demonstrated using succinate as substrate followed by inhibition of complex III with antimycin A (Fig. 6B,C). These substrate-specific measurements support the idea that electron flow from complexes I and II to ubiquinone/complex III is disrupted, indicating impaired coupling in the absence of mature CL in T cells. Notably, the enhanced OCR at complex I and the reduced activity at complex II suggest that CL differentially modulates the function of individual ETC complexes. These opposing – and potentially mutually neutralizing – effects may account for the lack of significant differences in the basal OCR observed in intact T cells. Additionally, compensatory mechanisms and the production of reactive oxygen species (ROS) may further ‘mask’ mitochondrial defects in the absence of Tafazzin when assessing OCR via extracellular flux analysis. Indeed, mitochondrial content per cell was increased in Tafazzin-deficient Th1 and Th17 cells (Fig. 6SF), suggesting that alterations in mitochondrial structure and/or number equalize impaired respiration efficiency. This compensatory response is further supported by the observation that relative membrane potential – calculated by the proton gradient at the IMM normalized to mitochondrial content – was significantly decreased in Tafazzin-deficient T cells (Fig. S6G).

**Figure 6.**
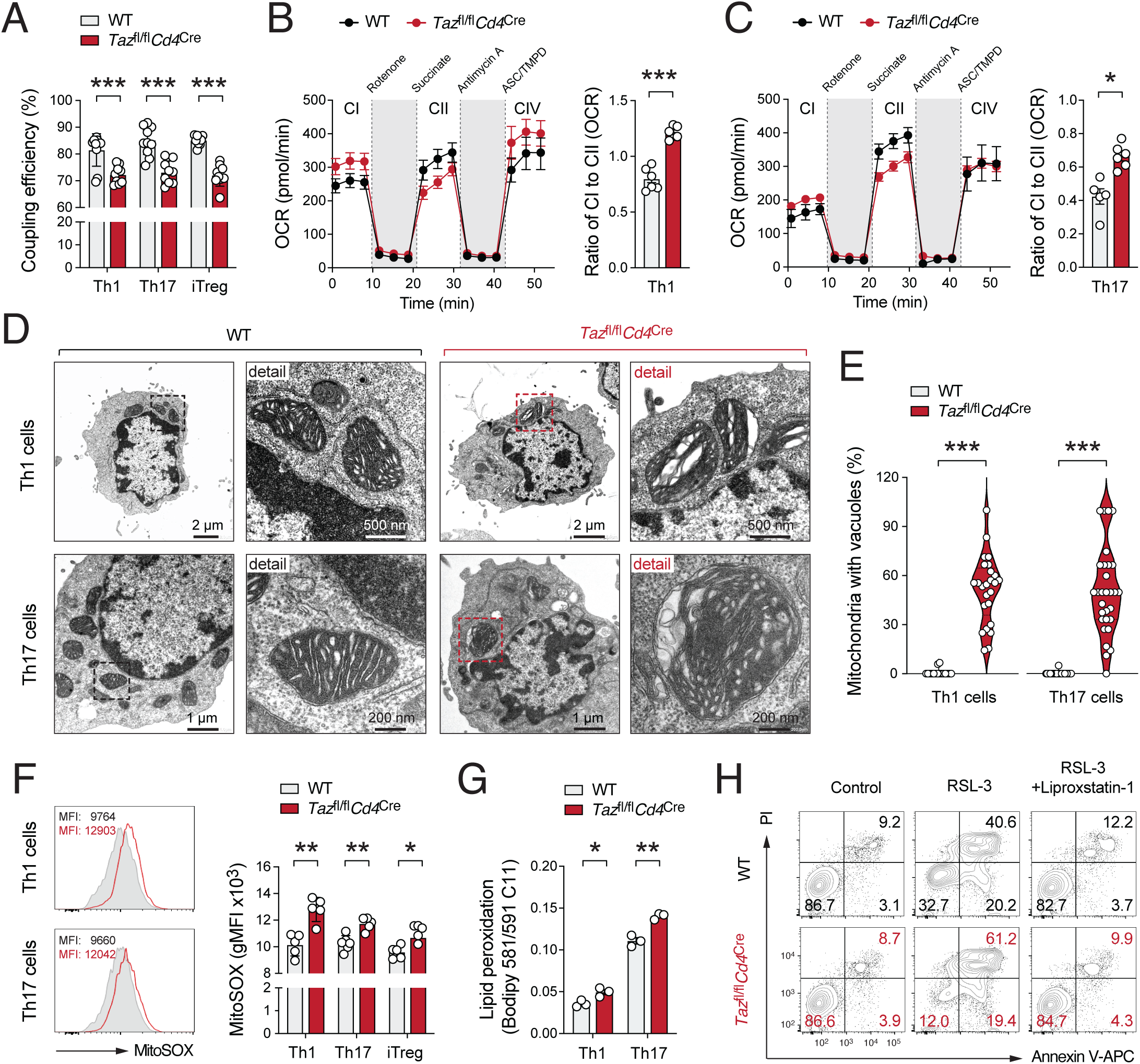
Cardiolipin remodeling controls cristae structure, ETC coupling efficiency and ROS production in T cells. **(A)** Analysis of electron transport chain (ETC) coupling efficiency in WT and Tafazzin-deficient Th1, Th17 and iTreg cells using a Seahorse extracellular flux analyzer; means ± SEM of 9-11 samples. Coupling efficiency was calculated by (ATP production rate) / (basal respiration rate) x 100; means ± SEM of 9-11 samples. (**B** and **C**) Electron flow assay at ETC complexes I, II and IV using a Seahorse extracellular flux analyzer in Th1 (B) and Th17 cells (C); means ± SEM of 5-6 samples. (**D**) Representative transmission electron microscopy (TEM) micrographs of mitochondrial cristae organization in WT and Tafazzin-deficient Th1 and Th17 cells, size is indicated by scale bars. **(E)** Frequency of mitochondria per cell with cristae delamination and/or vacuole-like structures as shown in (D); analysis of > 25 individual Th1 and Th17 cells from two independent experiments. **(F)** Flow cytometric analysis of mitochondrial reactive oxygen species (ROS) in WT and Tafazzin-deficient Th1, Th17 and iTreg cells using MitoSOX probes; means ± SEM of 5-6 samples. **(G)** Flow cytometric analysis of lipid peroxidation in WT and Tafazzin-deficient Th1 and Th17 cells using Bodipy 581/591 C11; means ± SEM of 3 mice. **(H)** Representative cell death analysis in WT and Tafazzin-deficient Th17 cells after treatment with the GPX4 inhibitor RSL-3 using flow cytometry. Statistical analyses in (A-C) and (E-G) by unpaired Student’s t-tests. *, p<0.05; **, p<0.01, ***, p<0.001.

Building on the observation that the bioenergetic coupling at the ETC complexes is impaired in the absence of Tafazzin-mediated CL remodeling, we examined mitochondrial morphology and cristae organization using transmission electron microscopy (TEM). Consistent with the elevated MitoTracker Red signal (Fig. 6SF), mitochondria in Tafazzin-deficient Th1 and Th17 cells appeared enlarged (Fig. S6H). Although the number of mitochondria per cell was unchanged or slightly decreased in our TEM analyses, Tafazzin-deficient mitochondria exhibited a’swollen’ morphology, characterized by pronounced cristae delamination and the presence of vacuole-like structures within the mitochondrial matrix (Fig. 6D). Notably, nearly all Tafazzin-deficient Th1 and Th17 cells displayed fragmented cristae, with 10-90% mitochondria per cells affected (Fig. 6E). Similar structural abnormalities have been previously reported in cardiomyocytes, fibroblasts and lymphoblasts from Tafazzin-deficient mice and BTHS patients, and are associated with impaired respiration and increased ROS production ^47,48^. In addition to stabilizing ETC complexes, preventing mitochondrial ROS production and facilitating their assembly into respirasomes, polyunsaturated CL species also serve as a “redox buffer” within the IMM by scavenging mitochondrial ROS through peroxidation of their double bounds, thereby preserving mitochondrial integrity and function ^8,48^. Furthermore, in absence of tafazzin, accumulated MLCL forms a peroxidase complex with cytochrome c that can oxidize polyunsaturated lipids ^49,50^. In line with this concept, Tafazzin-deficient T cells exhibited not only elevated mitochondrial ROS levels (Fig. 6F) but also increased lipid peroxidation (Fig. 6G).

The elevation of cellular ROS and lipid peroxidation are hallmarks of ferroptosis, a form of regulated, oxidative cell death. However, despite elevated mitochondrial ROS production and lipid peroxidation, Tafazzin-deficient T cells did not exhibit any significant reduction in viability under steady-state conditions (Fig. S2I and Fig. S3J). This resilience to redox stress is likely due to metabolic rewiring in these cells, particularly through the ISR pathway (Fig. 5C-H and Fig. S5G, H), which induce the expression of amino acid transporters (e.g., SLC1A4, SLC1A5, SLC7A11) and enzymes (ASNS, CTH, PYCR1) that promote glutathione biosynthesis, thereby mitigating oxidative stress (Fig. 5 and Fig. S6I). To test the hypothesis that elevated ISR protects against oxidative damage in the absence of CL, we inhibited glutathione peroxidase 4 (GPX4), a key regulator of lipid peroxidation and ferroptosis, using RSL-3. As expected, RSL-3 treatment resulted in increased lipid peroxidation and cell death of T cells (Fig. 6H and Fig. S6I-K), effects that could be rescued by the ferroptosis inhibitor liproxstatin-1. Importantly, Tafazzin-deficient Th17 cells were significantly more sensitive to RSL-3-induced oxidative damage compared to WT controls (Fig. 6H and Fig. S6I-K), suggesting that ISR-mediated metabolic rewiring and increased glutathione biosynthesis protect Tafazzin-deficient T cells from oxidative cell death under steady-state conditions.

Altogether, our findings underscore that Tafazzin-mediated CL remodeling is essential for optimal mitochondrial respiration in T cells. CL remodeling enhances chemiosmotic coupling efficiency, limits mitochondrial ROS production and supports cellular redox homeostasis, thereby promoting the metabolic resilience and effector functions of inflammatory T cells.

## DISCUSSION

T cell activation and effector differentiation are associated with significant structural and functional changes in the mitochondrial membranes. These changes also impact the composition of membrane lipids, including CL species ^6,10,11,18,51^. CL plays a critical role in facilitating supercomplex formation, enhancing respiratory efficiency and limiting ROS production ^10,51,52^. Consistent with this, our study demonstrates that the CL composition in CD4^+^ T cells is highly dynamic, with a decrease of MLCL and an accumulation of various CL species during Th cell differentiation. The composition and remodeling of CL vary in both acyl chain length and saturation. Tissues with high oxidative activity, such as cardiac muscle, tend to have elevated levels of CL species containing polyunsaturated side chains, such as CL(72:08), which features four linoleoyl acyl chains ^53^. However, CL composition differs across various tissues and cellular states, reflecting their specific metabolic and functional demands as well as acyl chains in the pool of available phospholipids ^53–57^. In agreement with previous studies ^34,36,58^, our findings revealed a broad and dynamic distribution of CL species in naïve and differentiated T cells. Notably, the CL to MLCL ratio increases upon T cell activation, suggesting that late-stage CL maturation plays a key role in mitochondrial remodeling during CD4^+^ T cell differentiation.

To directly investigate the role of Tafazzin-mediated CL remodeling on T cell function, we utilized the standard mouse model for BTHS with systemic, doxycycline-inducible knockdown of *Tafazzin* mRNA (*Taz*^KD^ mice) ^38,39^ as well as transgenic mice with T cell-specific ablation of the *Taz* gene ^43^ (*Taz*^fl/fl^*Cd4*^Cre^ mice). In both models, Tafazzin-mediated CL maturation was almost completely abolished in T cells, while MLCL species were markedly enriched, as expected ^10,11,59^. Given that mature CL species were nearly undetectable, our results suggest that other acyltransferases, such as ALCAT1, MLCLAT and the recently discovered PLAAT1 ^18,23,60^, are unable to compensate for the loss of Tafazzin in CD4^+^ T cells. Consequently, loss of CL remodeling in Tafazzin-deficient T cells resulted in marked ultrastructural changes, characterized by a ‘swollen’ mitochondrial phenotype, cristae disorganization and the appearance of vacuole-like structures within the mitochondrial matrix – features previously observed in BTHS patient-derived lymphoblasts ^47^. These morphological abnormalities were associated with inefficient coupling of the ETC complexes, diminished respiratory capacity and increased ROS production in Tafazzin-deficient Th1 and Th17 cells. To meet their bioenergetic demands, these cells increased the total mitochondrial content as a compensatory mechanism for the functional deficits of the individual organelles. While this adaption partially mitigated the OXPHOS impairment in the absence of CL, it failed to prevent elevated mitochondrial ROS production, lipid peroxidation and disruption of cellular redox homeostasis.

Integrated transcriptomic and metabolomic analyses revealed the molecular consequences of impaired mitochondrial function and elevated ROS levels in CL-deficient T cells. These analyses uncovered widespread metabolic alterations in Tafazzin-deficient T cells, affecting pathways such as fatty acid and cholesterol biosynthesis, amino acid metabolism and purine and pyrimidine metabolism. These pathophysiological metabolic adaptations observed in CD4^+^ T cells resemble those reported in other Tafazzin-deficient cell types, including cardiomyocytes, fibroblasts and neutrophils ^61,62^, suggesting a conserved metabolic rewiring mechanism that compensates for mitochondrial insufficiency in the absence of CL ^63^. Notably, we identified the integrated stress response (ISR) pathway as a key mediator linking mitochondrial dysfunction to the transcriptional and translational reprogramming of T cells. The ISR is a conserved cellular signaling pathway that enables adaption to various internal and external stressors, including ER stress and the unfolded protein response (UPR), nutrient deprivation and oxidative damage ^44,64^. In response to elevated mitochondrial ROS levels, the ISR is activated through phosphorylation of the translation initiation factor eIF2α, leading to global suppression of cap-dependent protein translation while promoting the expression and activation of ATF4 ^44,64^. In turn, ATF4 induces a pro-survival gene expression program to cope with amino acid deprivation and oxidative stress, thereby re-establishing cellular homeostasis ^65^. Among the ATF4-upregulated amino acid transporters in Tafazzin-deficient T cells were *Slc1a4* (ASCT1), *Slc1a5* (ASCT2) and *Slc7a11* (xCT), which are involved in glutathione biosynthesis to limit ROS accumulation, lipid peroxidation and cellular redox stress ^66^. Glutathione serves as a substrate for the selenoenzyme GPX4, which protects from lipid peroxidation and ferroptotic cell death ^67^. Consequently, inhibition of GPX4 exacerbated ROS-induced lipid peroxidation and ferroptosis in Tafazzin-deficient T cells. Although spontaneous ferroptosis was not observed, these findings suggest that under metabolically challenging conditions – such as amino acid deprivation within inflamed tissue niches – Tafazzin-deficient CD4^+^ T cells may be less resilient to oxidative stress. This vulnerability could account for the reduced numbers of Tafazzin-deficient Th1 and Th17 cells observed in inflamed tissues during autoimmune responses. Similar findings have recently been reported in cardiomyocytes and neutrophils, suggesting a conserved metabolic stress response in the absence of Tafazzin ^27,31,61,68,69^.

While upregulating ATF4 expression and stress response pathways, eIF2α simultaneously suppresses cap-dependent translation initiation, leading to a global reduction in mRNA translation ^44,64^. Although *Ifng* and *Il17* gene expression remained unchanged in Tafazzin-deficient Th1 and Th17 cells, the translation of cytokine mRNAs into protein was significantly impaired. The notion that the ISR impairs inflammatory cytokine production in absence of CL is further supported by the observation that treatment of Tafazzin-deficient Th1 and Th17 cells with an eIF2α inhibitor restored their cytokine expression. These findings are consistent with a recent study showing that the ISR regulates IFNγ and IL-17 production in tissue-resident T cells by modulating cytokine mRNA translation ^46^. Thus, adaption to metabolic stress in Tafazzin-deficient T cells has dual functional consequences: On the one hand, it mitigates oxidative stress to restore cellular homeostasis; on the other hand, it compromises the effector functions of inflammatory T cells, such as cytokine production. This transcriptional, translational and metabolic reprogramming of Tazazzin-deficient lymphocytes provides a plausible explanation for the alleviated T cell-mediated immunopathology in models of autoimmune encephalomyelitis and colitis.

Although impaired effector function and reduced metabolic resilience of Tafazzin-deficient Th cells confer protection against autoimmunity, they may also compromise protective immune responses to pathogens. This has potential relevance for BTHS patients, who commonly experience recurrent bacterial and fungal infections, including upper respiratory tract infections, chronic aphthous stomatitis, gingivitis, perianal dermatitis, and sepsis ^25,27^. While the increased susceptibility in BTHS has traditionally been attributed to persistent or intermittent neutropenia ^25,28^, our findings suggest that late-stage CL remodeling in Th1 and Th17 cells may contribute to the immunodeficiency observed in these patients. Similar to our observations in Tafazzin-deficient mice, the numbers of CD4^+^ and CD8^+^ T cells are generally normal in BTHS patients ^27,32^. However, whether defective CL remodeling also affects adaptive immunity in BTHS patients remains largely unknown, as only a limited number of studies have investigated the role of CL in T cells to date ^34,35,58,70,71^. Corrado and colleagues reported that PTPMT1-dependent CL biosynthesis is critical for effective cytotoxic CD8⁺ T cell responses to bacterial infections and for the development of pathogen-specific memory T cells ^34^. Similarly, Chen *et al.* demonstrated that PTPMT1-deficient T cells are unable to mount effective antitumor immunity^35^. However, it is important to note that PTPMT1 regulates the *de novo* biosynthesis of CL from phosphatidyl glycerophosphate, whereas Tafazzin mediates its late-stage remodeling, specifically the transacylation of MLCL to mature CL ^10,51^. Notably, Tafazzin deficiency did not recapitulate the functional defects observed in PTPMT1-deficient CD8^+^ T cells ^34^, suggesting that CL biosynthesis and remodeling impact CD8^+^ T cell differentiation and effector function through distinct mechanisms. Although Tafazzin seems dispensable for ‘cytotoxic’ T cell responses ^34^, our findings reveal a novel, cell-intrinsic role of Tafazzin-mediated CL remodeling in CD4^+^ T ‘helper’ cells. Whether CL fulfills a similar function in human lymphocytes, and whether defects in Th1 and Th17 cells contribute to the heightened susceptibility to infections observed in BTHS patients, remains to be investigated.

Collectively, our findings highlight novel and context-dependent roles of Tafazzin-mediated CL remodeling in shaping CD4^+^ T cell-mediated immune responses and underscore the need to further investigate its contributions to immune dysfunction in BTHS.

## ACKNOWLEDGEMENTS

We thank Drs. Douglas Strathdee (Cancer Research UK, the Beatson Institute, Glasgow, UK) for kindly providing *Taz*^fl/fl^ mice ^43^. We sincerely thank Dr. Vasco Sequeira (Comprehensive Heart Failure Center, Würzburg), Daniela Bunsen and Claudia Gehrig-Höhn (Imaging Core Facility of the Biocenter, Würzburg) for their support with animal experimentation and TEM analyses, respectively. This work was supported by the Deutsche Forschungsgemeinschaft (DFG) SFB-TR 124/3 (“FungiNet”) – project number: 210879364; SFB 1526/1 (“PANTAU”), project number: 454193335; SFB-TR 338/1 (“LETSimmun”) – project number: 452881907, SFB 1583/1 (“DECIDE”) – project number: 49262049; and individual project grants VA882/2-1 and VA882/3-2 (to M.V.). Further support was provided by the Barth Syndrome Foundation (to S.M.C.), the National Institutes of Health awards R01HL165729 and R01GM15174 (to S.M.C.), the SFB 1525/1 (“Cardio-Immune Interfaces”) – project number: 453989101 (to M.V., C.M. and J.D.) and the Mildred Scheel Early Career Center (MSNZ) of the German Cancer Aid (to M.V. and M.E.). C.M. was supported by DFG project numbers 505805397 and 315254108. The JEOL JEM1400 electron microscope is funded by the DFG – 426173797 (INST 93/1003-1 FUGG).

## AUTHOR CONTRIBUTION

Participated in research design: M.V., X.Z., W.S., C.M., C.S., S.M.C. and J.D. Conducted experiments: X.Z. S.H., M.H., H.W., K.S., K.J.E., W.S., M.E., S.L.P. and W.S. Performed data analysis: M.V., S.M.H., M.E., C.S., H.W. and W.S. Wrote the manuscript: M.V. and X.Z.

## DECLARATION OF INTERRSTS

The authors declare no competing interests.

## METHODS

### Animals

All mice were bred and maintained under specific pathogen free conditions in the Center for Experimental Medicine (ZEMM) or the Institute for Systems Immunology at the Julius-Maximilians University of Würzburg. Mice were maintained on a 12/12 h light/dark cycle at between 20-24 °C in individually ventilated cages. Mice had access to standard chow (Ssniff; cat# V1534) and autoclaved water *ad libitum* and health status of the animals was inspected daily by the responsible caretakers. Hygiene status of the sentinel mice was monitored quarterly according to the FELASA guidelines. Both male and female mice between 2 and 8 months of age at the time of the experiment were used for the *in vitro* experiments described in this study. Respective animal protocols were approved by government of Lower Franconia, Germany. C57BL/6 (JAX strain 000664), CD45.1^+^ (strain 002014), *Rosa26^H1/tetO-shRNA:TAZ^*(strain 014648), *Rag1^-/-^* (strain 002216) and *Cd4*^Cre^ mice (strain 017336) were purchased from the Jackson Laboratories (JAX) and/or maintained in our institution. *Taz*^fl/fl^ mice ^43^ were generated and kindly provided by Douglas Strathdee (Cancer Research UK, the Beatson Institute, Glasgow, UK). All animals used in this study were on a C57BL/6J genetic background.

### Knockdown of Tafazzin in mice

Systemic knockdown of *Tafazzin* in *ROSA26^H1/tetO-shRNA:TAZ^* (*Taz*^KD^) mice was previously described ^38^. In brief, doxycycline (625 mg/kg) was administered as part of the standard rodent chow to control and *Taz*^KD^ animals. To avoid male infertility during breeding, female mice were treated with doxycycline for one week and doxycycline was withdrawn during mating. After detecting copulatory plugs, Doxycycline treatment was continued.

### T cell activation and differentiation

For *in vitro* cultures, murine CD4^+^ T cells were isolated from single cell suspension of lymph nodes and spleen by negative selection using the MojoSort Mouse CD4 T cell isolation kit (BioLegend). T cells were cultured in modified RPMI 1640 medium with physiological glucose concentration (i.e., 100 mg/dL) by diluting standard RPMI 1640 medium (Gibco) with glucose-free RPMI medium (Roth). The medium was supplemented with 10% FBS (Sigma), 50 μM 2-mercaptoethanol (b-ME), 1% penicillin/streptomycin and 1% GlutaMAX-I (all Gibco) and the cells were cultured at 37°C with 5% CO2. For T cell differentiation, delta-surface plates (Nunc) were pre-coated with 12 μg/ml polyclonal anti-hamster IgG (MP Biomedicals) for 2 h and washed once with PBS. In 24-well plates, 1 x10^6^ cells were activated with 0.25 μg/ml (for Th17) or 0.5 μg/ml (for Th1 and iTreg subsets) of anti-CD3 (clone 145-2C1) together with 1 μg/ml anti-CD28 (clone 37.51, both BioXCell) and polarized into different Th subsets using the following cytokines and antibodies: For Th1 cells: 2.5 μg/ml anti-IL-4 (clone 11B11), 10 ng/ml rhIL-2 and 10 ng/ml rmIL-12; for Th17 cells: 2.5 μg/ml anti-IL-4 (clone 11B11), 2.5 μg/ml anti-IFNγ (clone XMG1.2), 20 ng/ml rmIL-6 and 0.5 ng/ml rhTGFb1 and for iTreg cells: 2.5 μg/ml anti-IL-4 (clone 11B11), 2.5 μg/ml anti-IFNg (clone XMG1.2), 10 ng/ml rhIL-2 and 5 ng/ml rhTGFb1. All cell culture antibodies and recombinant cytokines were from BioXCell and Peprotech, respectively. To induce ISR during Th cell differentiation, CD4^+^ T cells were activated for 24h with anti-CD3/CD28 and treated with 0.5 or 1 µM NaAsO2 (Sigma-Aldrich) for 48h. To inhibit the ISR pathway, T cells were incubated with 500 nM ISRIB (MedChemExpress) after 24h anti-CD3/CD28 stimulation for two days.

### Flow cytometry

Flow cytometric staining was performed as previously described ^72^. Briefly, cells were stained with Fixable Viability Dye eFluor 780 (eBioscience) for 10 min in PBS at RT together with an anti-FcgRII/FcgRIII antibody (clone 2.4G2; Bio X Cell) to prevent unspecific binding. After washing, surface antigens were stained with fluorophore-conjugated antibodies in PBS containing 0.5% BSA for 20 min at RT in the dark. For intracellular cytokine staining, cells were stimulated with 1 μM ionomycin (BioMol) and 30 nM phorbol-12-myristat-13-acetate (PMA, Sigma) in the presence of 2 μg/ml brefeldin A (eBioscience) for 5 h at 37°C. After surface staining, cells were fixed with IC-fixation buffer (eBioscience) and intracellular cytokines were stained using 1x permeabilization buffer (both eBioscience) at RT. For detection of intranuclear antigens, cells were fixed using Foxp3/TF staining buffer set according to manufacturer’s recommendations (eBioscience). To analyze T cell proliferation, CD4^+^ T cells were loaded with 5 μM CellTrace Violet (Invitrogen) according to manufacturer’s instructions. Glucose uptake by T cells *in vitro* was monitored using the fluorescent glucose analog 2-NBDG (Cayman Chemicals). After starving for 15 min in glucose-free medium, T cells were incubated with 300 μM 2-NBDG for 30 min at 37°C, washed and processed for flow cytometric analysis. Neutral lipid content of cultivated Th cells was measured using the BODIPY 493/503 reagent (Invitrogen). For the quantification of the mitochondrial volume, membrane potential and ROS production, T cells were loaded in complete RPMI medium at 37°C with 500 nM MitoTracker deep red, 2 nM Tetramethylrhodamin-Ethylester (TMRE) and 2.5 µM MitoSOX, respectively (all Invitrogen). As background control for TMRE, T cells were pre-treated with 20 µM Trifluoromethoxy carbonylcyanide phenylhydrazone (FCCP, Cayman Chemicals) for 15 min to depolarize the membrane potential. To measure the phospho-eIF2α levels, T cells were subsequently fixed with IC-fixation buffer (BioLegend) for 40 min at RT, permeabilized and stained with the primary antibody against phospho-eIF2α (1:1000, Invitrogen) for 40 min at RT. After two rounds of washing, an anti-rabbit secondary antibody conjugated with AF647 against Rabbit IgG (1:1000, Invitrogen) was used to detect phosphorylated eIF2α. All sample acquisition was performed with a BD Celesta flow cytometer (BD Biosciences) or an Aurora Flow Cytometer (Cytek) and further analyzed with the FlowJo software (Tree Star). The gating strategies for flow cytometric analyses can be found in Fig. S7. A complete list of antibodies, clones and conjugates used in this study can be found in Table S1.

### Seahorse extracellular flux analysis

Mitochondrial respiration and lactate secretion of T cells was measured as their oxygen consumption rate (OCR) and glycolytic proton efflux rate (glycoPER), respectively, using an oxygen-controlled XFe96 extracellular flux analyzer (Seahorse Bioscience). XFe96 cell culture microplates (Agilent) were pre-coated with 22 μg/ml Cell-Tak (Corning) and 2×10^5^ T cells per well were attached in 5-8 replicates in Seahorse XF RPMI medium (Agilent) supplemented with 2 mM L-glutamine (Gibco), 1 mM sodium pyruvate (Sigma) and 10 mM D-glucose (Sigma). After incubation for 1 h in a CO_2_-free incubator at 37°C, glycolytic and mitochondrial stress tests were performed. For assessing glycolysis, basal extracellular acidification rate (ECAR) was measured followed by addition of 0.5 μM rotenone (AdipoGen) and 0.5 μM antimycin A (Sigma) to inhibit mitochondrial complex I and III, respectively. At the end of the measurement, 50 mM 2-DG (Sigma) was added to completely block glycolysis. To analyze mitochondrial respiration, basal oxygen consumption was measured followed by the addition of 2 μM oligomycin (Cayman Chemicals), an ATP synthase inhibitor, 1 μM of the protonophore carbonyl cyanide-*4*-(trifluoromethoxy)-phenylhydrazone (FCCP, Cayman Chemical) and 0.5 μM rotenone (AdipoGen) together with 0.5 μM antimycin A (Sigma). The basal oxygen consumption was calculated by subtracting the OCR after rotenone and antimycin A treatment from the OCR before oligomycin treatment. The maximal OCR was calculated by subtracting the OCR after rotenone and antimycin A treatment from the OCR measured after addition of FCCP. To perform an electron flow assay using the Seahorse XFe96, 2×10^5^ differentiated CD4^+^ T cells per well were seeded in mitochondrial assay solution (Agilent) supplemented with 10 mM sodium pyruvate (Sigma), 1 mM malate (Carl Roth), 4 mM ADP (Sigma) and 1 nM Agilent Seahorse XF Plasma Membrane Permeabilizer (PMP). The following compounds were injected into the culture plate sequentially: 1 µM rotenone (Biomol) to inhibit mitochondrial complex I, 10 mM sodium succinate (BioLegend) as a substrate for complex II, 2 µM antimycin A (Sigma) to inhibit complex III, and a mixture of 10 mM Ascorbate (Sigma) and 100 µM TMPD (Sigma) as substrates for complex IV.

### Metabolomic profiling

Metabolite extraction: Snap-frozen pellets of > 1×10^7^ T cells were resuspended in 360 µl 10 mM HCl/MeOH (17/19, v/v) containing external water-soluble standard compounds (0.02 µM lamivudine and 2 µM each of D_2_-glucose, D_4_-succinate, D_5_-glycine and ^15^N-glutamate) and treated with ultrasound (Branson). External lipid standard compounds (100 µM D_7_-cholesterol, 20 µM D_31_-palmitic acid and 10 µM each of LPA-(17:0) and LPC-(17:0)) in 20 µl MeOH/CHCl_3_ and 90 µl CHCl_3_ were added. After vigorous mixing, another 100 µl CHCl_3_ were added, and the resulting phases separated after mixing and centrifugation. The lower phases were re-extracted with 150 µl synth. upper phase (CHCl_3_/MeOH/H_2_O: 4.1/47.3/48.8, v/v/v) and the combined upper phases were re-extracted with 300 µl synth. lower phase (CHCl_3_/MeOH/H_2_O: 58.4/33.4/8.0, v/v/v). The combined lower phases were evaporated to dryness under a stream of nitrogen, the combined upper phases were evaporated under a stream of nitrogen for 15 min and taken to dryness in a centrifugal evaporator. Liquid chromatography and mass spectrometry (LC/MS) analyses were performed on a Dionex Ultimate 3000 UHPLC system connected to a Q Exactive mass spectrometer (QE-MS) equipped with a HESI probe (both Thermo Scientific).

Water soluble metabolites: Upper phase residues were reconstituted in 100 µl of 5 mM NH_4_OAc in CH_3_CN/H_2_O (50/50, v/v). Chromatographic separation was achieved by applying 3 µl sample on a XBridge Premier BEH Amide (100 × 2.1 mm, 2.5 μm) (Waters), protected by a Supelco ColumnSaver particle filter (Merck) and a gradient of mobile phase A (5 mM NH_4_OAc in CH_3_CN/H_2_O: 40/60, v/v) and mobile phase B (5 mM NH_4_OAc in CH_3_CN/H_2_O: 95/5, v/v) maintaining a flow rate of 200 µl/min. The LC gradient program was 100% mobile phase B for 2 min, followed by a linear decrease to 20% B within 23 min, maintaining 20% B for 21 min and returning to 100% B in 2 min, followed by 7 min 100% B for column equilibration before each injection. The eluent was directed to the QE-MS from 2.7 min to 46 min after sample application. Mass detection was conducted in alternating pos/neg full scan mode (at 70k resolution, scan range m/z 69 - 1000, AGC target 1E6 and 200 ms max. injection time). HESI parameters: Sheath gas: 20, aux gas: 1, spray voltage: 3.0 kV, capillary temp.: 300 °C, S-lens RF level: 50.0, aux gas heater temperature: 120 °C.

Lipid metabolites: Lower phase residues (lipid fraction of the B/D extraction) were reconstituted in 100 µl of iPrOH. Lipid samples were measured twice, applying hydrophilic interaction liquid chromatography (HILIC) and reverse phase (RP) conditions. HILIC chromatographic separation was achieved by applying 3 µl sample on a XBridge Premier BEH Amide (100 x 2.1 mm, 2.5 μm) (Waters), protected by a Supelco ColumnSaver particle filter (Merck) and a gradient of mobile phase A (5 mM NH_4_OAc in CH_3_CN/H_2_O: 40/60, v/v) and mobile phase B (5 mM NH_4_OAc in CH_3_CN/H_2_O: 95/5, v/v) maintaining a flow rate of 200 µl/min. The LC gradient program was 100% mobile phase B for 2 min, followed by a linear decrease to 20% B within 18 min, maintaining 20% B for 10 min and returning to 100% B in 2 min, followed by 7 min 100% B for column equilibration before each injection. The eluent was directed to the QE-MS from 2.7 min to 16 min after sample application. RP chromatographic separation was achieved by applying 3 µl sample on a Acclaim C8 column (100 x 2.1 mm, 3 μm) (Thermo Scientific), protected by a Supelco ColumnSaver particle filter (Merck) and a gradient of mobile phase A (MeOH/H_2_O/FA: 5/94.9/0.1, v/v/v), mobile phase B (iPrOH/CH_3_CN/H_2_O/FA: 45/45/9.9/0.1, v/v/v/v) and mobile phase C (5 mM NH_4_OAc in CH_3_CN/H_2_O: 95/5, v/v) maintaining a flow rate of 200 µl/min. The LC gradient program was 20% mobile phase B for 2 min, followed by a linear increase to 100% B within 10 min, maintaining 100% B for 20 min, switching to 100% C for 16 min and returning to 20% B in 2 min, followed by 7 min 20% B for column equilibration before each injection. The eluent was directed to the QE-MS from 4.9 min to 48 min after sample application. Mass detection was conducted in alternating pos./neg. full scan mode (at 70k resolution, scan range m/z 190 - 1400, AGC target 1E6 and 200 ms max. injection time). HESI parameters: sheath gas: 20, aux gas: 1, spray voltage: 3.0 kV, capillary temp.: 300 °C, S-lens RF level: 50.0, aux gas heater temperature: 120 °C. Manual curation and integration of chromatographic peaks were performed with TraceFinder 5.1 using a mass tolerance of ± 2 mMUs. Metabolite set enrichment analysis (MSEA) and combined network analyses of differential metabolites and gene expression were performed with the MetaboAnalyst software (https://www.metaboanalyst.ca).

### qRT-PCR and RNA sequencing

Total RNA was extracted using the Roti-Prep Mini Kit (Roth) and cDNA was synthesized using the iScript cDNA synthesis kit (Bio-Rad). Quantitative real-time PCR was performed using the SYBR Green qPCR Master Mix (Bio-Rad) and specific primers (Table S2). The relative abundance of transcripts was normalized to the expression of housekeeping genes (18S) using the 2^-ΔCt^ method. For bulk RNA-sequencing (RNA-seq), naïve T cells of *Taz*^fl/fl^*Cd4*^Cre^ or WT mice were differentiated for 3 days into Th1, Th17 and iTreg cells and directly harvested in RNAprotect Cell reagent and stored at-80°C o/n before total RNA was extracted using the RNeasy Plus Micro Kit (both Quiagen). RNA quality and quantity were checked using a 2100 Bioanalyzer with the RNA 6000 Nano kit (Agilent Technologies). The RIN for all samples was > 8. cDNA libraries for sequencing were prepared from 500 ng of total RNA with TruSeq mRNA Stranded Library Prep Kit from Illumina according to manufacturer’s instructions (1/2 volume). Libraries were quantified by QubitTM 3.0 Fluometer (Thermo Scientific) and quality was checked using 2100 Bioanalyzer (Agilent) with High Sensitivity DNA kit (Agilent). Libraries were quantified again by QubitTM 3.0 Fluometer (Thermo Scientific) and quality was checked using 2100 Bioanalyzer with High Sensitivity DNA kit (Agilent) before pooling.

RNA sequencing of pooled libraries, spiked with 1% PhiX control library, was performed at 19-36 million reads/sample in single-end mode with 75 nt read length on the NextSeq 500 platform (Illumina) with 1 High Output Kit v2.5. Demultiplexed FASTQ files were generated with bcl2fastq2 v2.20.0.422 (Illumina). To assure high sequence quality, Illumina reads were quality-and adapter-trimmed via Cutadapt version 2.5 using a cutoff Phred score of 20 in NextSeq mode and reads without any remaining bases were discarded. Processed reads were subsequently mapped to the mouse genome (GRCm38.p6 primary assembly and mitochondrion) using STAR v2.7.2b with default parameters based on RefSeq annotation version 108.20200622 for GRCm38.p6. Read counts on exon level summarized for each gene were generated using featureCounts v1.6.4 from the Subread package. Multi-mapping and multi-overlapping reads were counted non-strand-specific with a fractional count for each alignment and overlapping feature. The count output was utilized to identify differentially expressed genes using DESeq2 version 1.24.0. Read counts were normalized by DESeq2 and fold-change shrinkage was applied. Differences in gene expression were considered significant if padj < 0.01. For overlapping gene expression analyses, an online tool for comparing DEG lists with Venn’s diagrams (https://bioinfogp.cnb.csic.es/tools/venny/index.html) was used. For KEGG pathway analysis (https://www.bioinformatics.com.cn) and gene set enrichment analysis (GSEA, https://www.gsea-msigdb.org/gsea) public online platforms were used.

### Immunoblotting

Cells for immunoblotting were lysed in RIPA buffer (SantaCruz) supplemented with complete protease-inhibitor cocktail (Roche). Protein quantification was performed with the Pierce 660 nm Protein Assay (Pierce). Protein samples (40 µg) were separated via SDS-PAGE, blotted onto nitrocellulose membranes, blocked with 5% BSA and incubated with primary antibodies (murine anti-Tafazzin antibody clone 2G3F7 ^73^ was generated by Steven M. Claypool and the OXPHOS rodent antibody cocktail; Thermo Scientific). Detection was carried out using goat anti-mouse or anti-rabbit horseradish peroxidase conjugated secondary antibodies (BioRad) and visualized with the enhanced chemiluminescent SuperSignal reagent (Pierce) using the ChemiDoc imaging system (BioRad).

### Ferroptosis induction assay

To induce ferroptosis by GPX4 inhibition, 1.5×10^5^ CD4^+^ T cells per well were seeded in 96-well plates. After 24-hour activation with anti-CD3/CD28, 500 nM GPX4 inhibitor RSL-3 (Cayman Chemical) was used to induce ferroptotic cell death. As a control, 100 nM Liproxtatin-1 (Sigma) was added in some wells to inhibit RSL-3-induced lipid peroxidation and ferroptosis. Cell death was analyzed 48h later by flow cytometry using Annexin V and propidium iodide (both BioLegend). Lipid peroxidation and mitochondrial ROS levels were detected using Bodipy 581/591 C11 and MitoSOX reagents, respectively (both Thermo Scientific).

### Transmission electron microscopy

To analyze mitochondrial morphology in WT and Tafazzin-deficient Th and Th17 cell samples by transmission electron microscopy (TEM), 1×10^7^ differentiated CD4^+^ T cells were collected and washed once with pre-warmed PBS. Samples were prepared for TEM as previously described ^74^. The cell pellets were fixed with 2.5% glutaraldehyde and 100 mM sodium cacodylate (both Sigma) in ddH_2_O for 60 min, followed by incubation with 2% OsO_4_ (Sigma) for 90 min and o/n contrasting with 0.5% uranyl acetate. The cell pellets were dehydrated, embedded in Epon 812 (Sigma) and sliced into ∼ 80 nm ultrathin samples. TEM analysis was performed at 120 kV acceleration voltage at a JEM-1400 Flash transmission electron microscope (JOEL) equipped with a Matataki camera system. The number, size and morphology of mitochondria were analyzed using ImageJ (Fiji software).

### Experimental autoimmune encephalomyelitis (EAE)

To induce EAE in mice, 2 mg/ml MOG_35-55_ peptide (Synpeptide) was emulsified in IFA (BD) supplemented with 5 mg/ml *M. tuberculosis* H37Ra (Fisher Scientific) by syringe extrusion. In total 200 mg MOG_35-55_ peptide was subcutaneously (s.c.) injected into two different sites on the lower flanks of the mice, followed by intraperitoneal injection of 250 ng pertussis toxin (Enzo) on day 0 and 2. Mice were monitored daily for clinical signs of EAE and weight loss. The scores were assigned as described before ^75^. In brief, score 0: no paralysis, score 0,5: partially limp tail, score 1: paralyzed tail, score 2: uncoordinated movement, score 2,5: paralysis of one hind limb, score 3: paralysis of both hind limbs, score 3,5: weakness in forelimbs, score 4: paralysis of forelimbs and score 5: moribund. At the end of the experiment, mice were sacrificed and cell suspensions from the spleen, LNs and the spinal cords were prepared. Briefly, spinal cords were isolated, mined into small pieces and digested with 1 mg/ml collagenase D (Roche) and 20 μg/ml DNase I (Thermo Scientific) for 40 min at 37°C. After filtering through a 70 μm cell strainer, CNS-infiltrating lymphocytes were isolated by percoll gradient (30:70) centrifugation.

### Adoptive transfer colitis

For the induction of colitis, naïve CD4^+^CD25^-^CD45RB^high^ T cells were isolated from the spleens of WT and *Taz*^fl/fl^ *Cd4*^Cre^ littermate mice using a BD FACSAria III cell sorter as previously described ^76^. 5×10^5^ FACS-sorted T cells were transferred intraperitoneally into lymphopenic *Rag1*^-/-^ recipient mice and initiation of autoimmune colitis was monitored over the course of 8 weeks by analyzing weight loss of the recipient animals, stool samples and clinical signs of colonic inflammation, such as rectal prolapses. At the end of the experiment, mice were sacrificed before spleen, mesenteric LNs, small intestine and colon were extracted. For the isolation of lamina propria lymphocytes from the small intestine and the colon, tissues were washed with PBS and the intraepithelial fraction was dissociated by adding twice Ca^2+^ and Mg^2+^-free HBSS (Sigma) supplemented with 5 mM EDTA (Thermo Scientific) and 10 mM HEPES (Gibco) for 20 min at 37°C shaking. After rinsing with PBS, both tissues were minced into small pieces and digested with HBSS containing 500 μg/ml collagenase D (Roche), 20 μg/ml DNase I (Thermo Scientific) and 0.5 U/ml Dispase (BD) for 45 min at 37 °C shaking. Samples were filtered through 70 μm cell strainers and lymphocyte fractions were concentrated by percoll gradient (40% *vs.* 80%) centrifugation.

### Quantification and statistical analysis

The results are shown as mean ± standard error of the means (SEM). To determine the statistical significance of the differences between the experimental groups unpaired Student’s t or Mann-Whitney-U-tests tests were performed using the Prism 9 software (GraphPad). Sample sizes were based on experience and experimental complexity. Differences reached significance with p values < 0.05 (noted in figures as *), p < 0.01 (**), and p < 0.001 (***). The figure legends contain the number of independent samples or mice per group and the statistical analyses that were used.

### Data and code availability

The RNA sequencing (RNA-seq) dataset produced in this study is deposited under accession number GSE295044 (reviewer token: ydodgecgzhanzuf)

## SUPPLEMETARY INFORMATION

**Figure S1.**
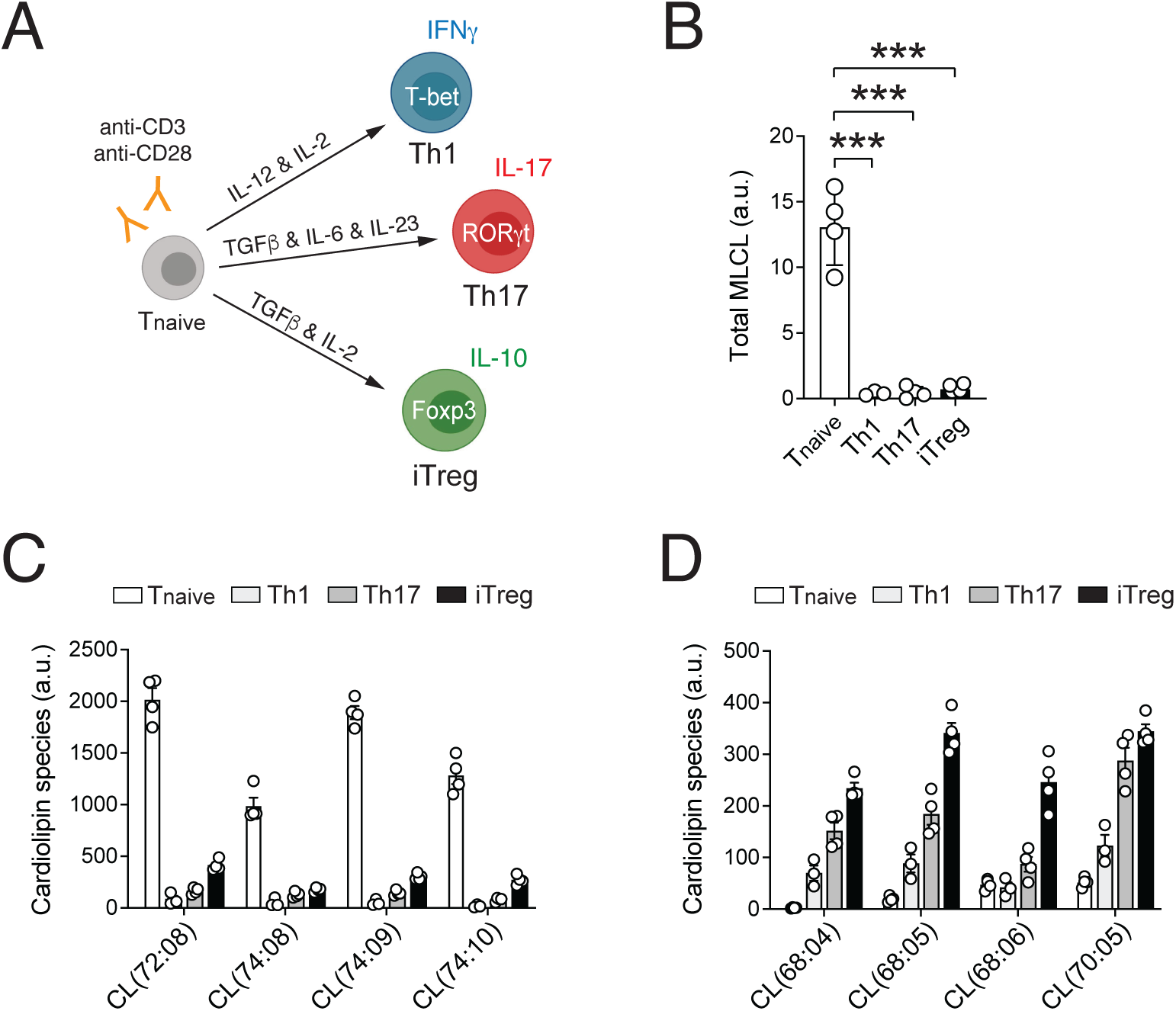
Cardiolipin composition in CD4^+^ T cell subsets. **(A)** Experimental design of Th1, Th17 and iTreg cell differentiation from naïve CD4^+^ T cells *in vitro*. **(B)** Analysis of total monolysocardiolipin (MLCL) levels per 1×10^6^ cells in naïve CD4^+^ T cells and after differentiation into Th1, Th17 and iTreg cells by liquid chromatography and mass spectrometry (LC/MS); means ± SEM of 4 biological replicates per group. **(C** and **D)** Most significantly decreased (C) and increased (D) cardiolipin (CL) species in naïve *versus* differentiated Th1, Th17 and iTreg cells analyzed by LC/MS; means ± SEM of 4 biological replicates per group. Statistical analyses in (B) by unpaired Student’s t-tests. ***, p<0.001.

**Figure S2.**
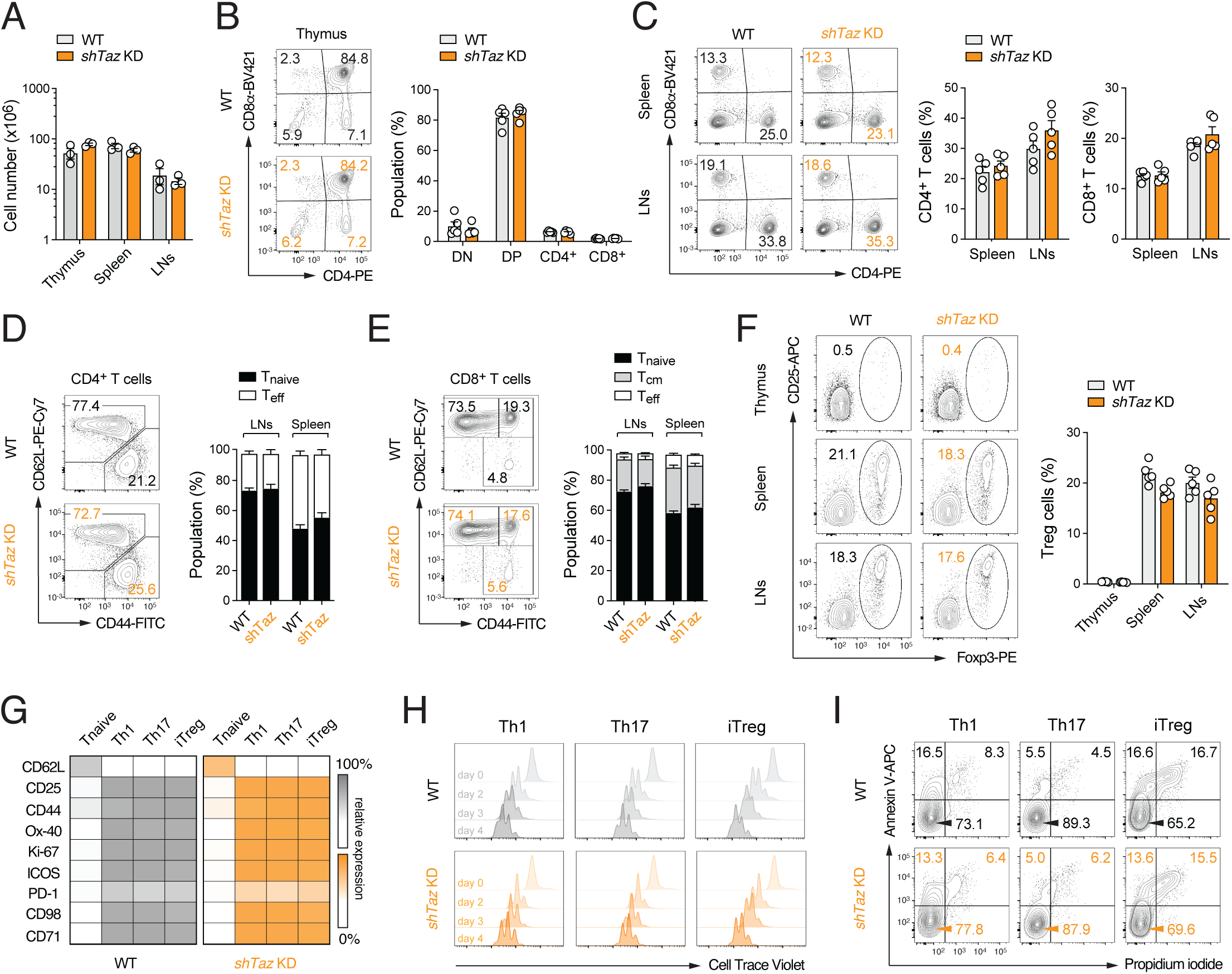
Systemic Tafazzin knockdown does not impair T cell development and activation. **(A)** Total cell numbers of thymus, spleen and lymph nodes (LNs) of WT and *Rosa26^H1/tetO-shRNA:TAZ^*(*Taz*^KD^) mice after doxycycline treatment for < 6 months; means ± SEM of 2-3 mice. **(B)** Flow cytometric analysis of thymic T cell development in WT and *Taz*^KD^ mice after doxycycline treatment for 6-8 months; means ± SEM of 5 mice. **(C)** Flow cytometric analysis of CD4^+^ and CD8^+^ T cells in spleens and LNs of WT and *Taz*^KD^ mice; means ± SEM of 5 mice. **(D** and **E)** Distribution of CD44^-^CD62L^+^ (naïve), CD44^+^ CD62L^+^ (central memory) and CD44^+^CD62L^-^ (effector) CD4^+^ (D) and CD8^+^ (E) T cells in WT and *Taz*^KD^ mice after doxycycline treatment for < 6 months; means ± SEM of 5 mice. **(F)** Analysis of Foxp3^+^ Treg cells in thymus, spleen and LNs of WT and *Taz*^KD^ mice after doxycycline treatment by flow cytometry; means ± SEM of 5 mice. **(G-F)** Systemic knockdown of Tafazzin does not impair T cell activation, proliferation and apoptosis *in vitro*. Representative flow cytometric analyses of selected activation marker expression (G), proliferation (H) and apoptosis in WT and *Taz*^KD^ T cells after *in vitro* differentiation into Th1, Th17 and iTregs cells.

**Figure S3.**
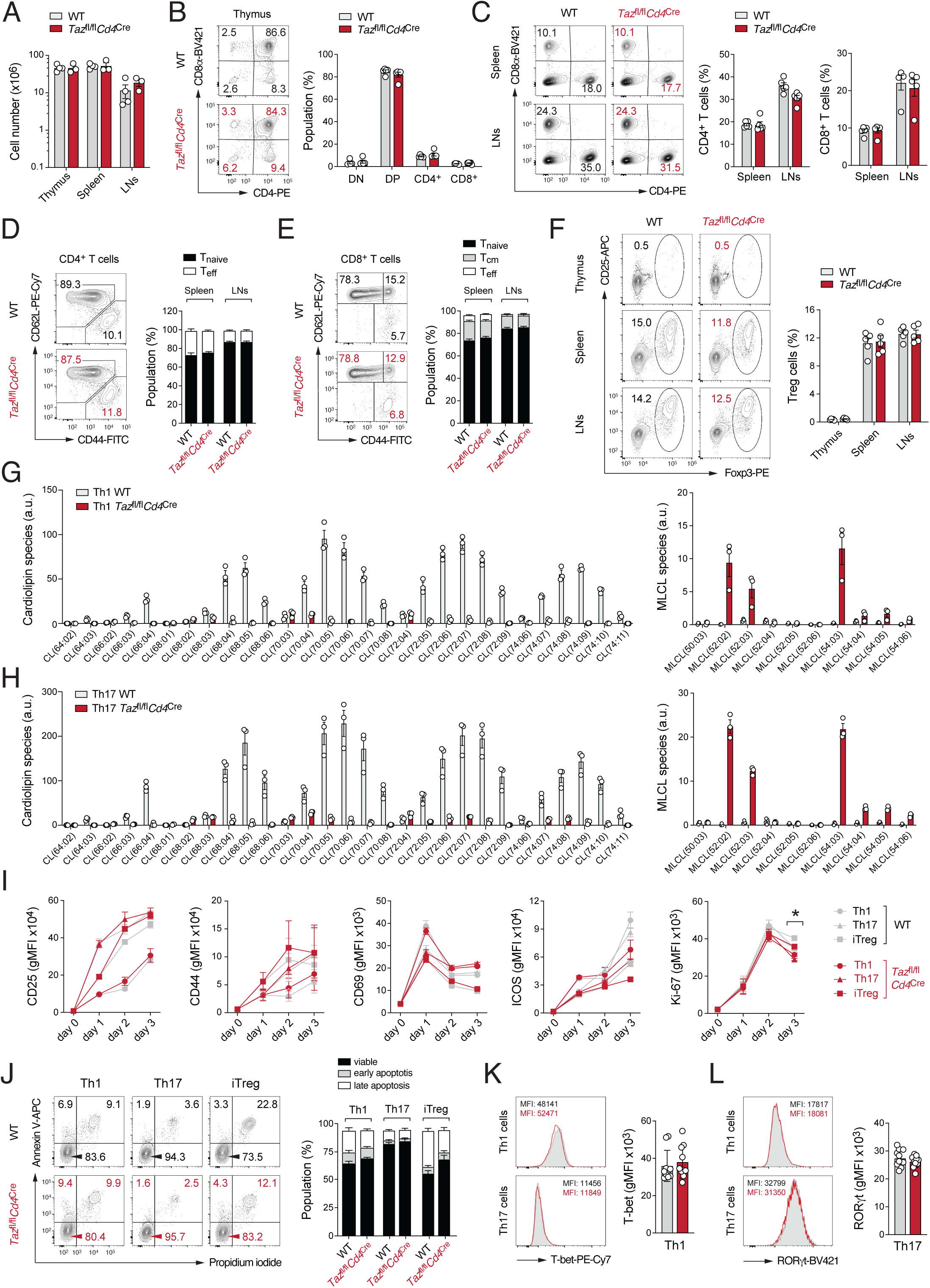
Characterization of mice with T cell-specific ablation of Tafazzin. **(A)** Total cell numbers of thymus, spleen and lymph nodes (LNs) of WT and *Taz*^fl/fl^*Cd4*^Cre^ mice; means ± SEM of 3-4 mice. **(B)** Flow cytometric analysis of thymic T cell development in WT and *Taz*^fl/fl^*Cd4*^Cre^ mice; means ± SEM of 3-4 mice. **(C)** Analysis of CD4^+^ and CD8^+^ T cells in spleens and LNs of WT and *Taz*^fl/fl^*Cd4*^Cre^ mice by flow cytometry; means ± SEM of 4-5 mice. **(D** and **E)** Distribution of CD44^-^CD62L^+^ (naïve), CD44^+^ CD62L^+^ (central memory) and CD44^+^CD62L^-^ (effector) CD4^+^ (D) and CD8^+^ (E) T cells in WT and *Taz*^fl/fl^*Cd4*^Cre^mice; means ± SEM of 4-5 mice. **(F)** Flow cytometric analysis of Foxp3^+^ Treg cells in thymus, spleen and LNs of WT and *Taz*^fl/fl^*Cd4*^Cre^mice; means ± SEM of 4-5 mice. **(G** and **H)** Analysis of cardiolipin (CL) and monolysocardiolipin (MLCL) species per 1×10^6^ cells in *in vitro* differentiated WT and Tafazzin-deficient Th1 (G) and Th17 cells (H) by liquid chromatography and mass spectrometry (LC/MS); means ± SEM 4 biological replicates per group. Numbers in brackets indicate number of carbon atoms to unsaturated C=C bonds of the acyl chains. **(I)** Flow cytometric analysis of CD25, CD44, CD69, ICOS and Ki-67 protein expression in WT and Tafazzin-deficient T cells during Th1, Th17 and iTreg differentiation; means ± SEM 5 mice. **(J)** Analysis of apoptosis in WT and Tafazzin-deficient Th1, Th17 and iTregs cells; means ± SEM 5 mice. **(K** and **L)** Analysis of T-bet (K) and RORγt (L) protein expression in Th1 and Th17 cells, respectively; means ± SEM 10-11 mice. Statistical analyses in (I) by unpaired Student’s t-tests. *, p<0.05.

**Figure S4.**
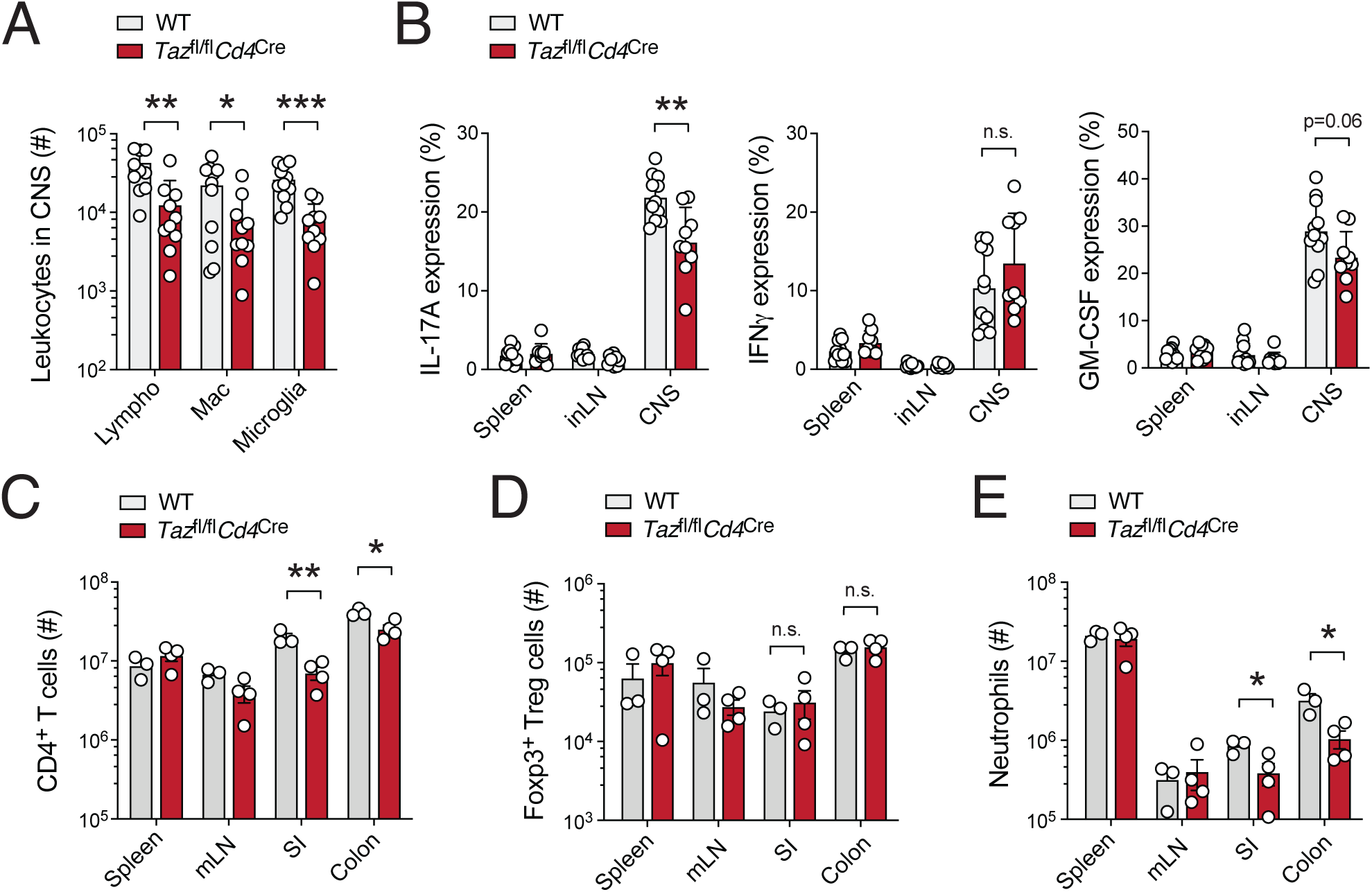
Ablation of Tafazzin in T cells alleviates autoimmunity in mice. **(A)** Analysis of lymphocyte, macrophage and microglia cell numbers in the CNS of WT and *Taz*^fl/fl^*Cd4*^Cre^ mice 18 days after immunization with MOG_35-55_ peptide emulsified in CFA; means ± SEM of 10-11 mice per cohort. **(B)** Frequencies of IL-17A, IFNγ and GM-CSF producing CD4^+^ T cells in the spleen, inguinal LNs and CNS of WT and *Taz*^fl/fl^*Cd4*^Cre^ mice 18 days after immunization with MOG_35-55_ peptide and restimulation with PMA/Iono for 5 h; means ± SEM of 10-11 mice. **(C-E)** Tafazzin-deficient T cells fail to induce autoimmune colitis after adoptive transfer into lymphopenic (*Rag1*^-/-^) mice. Absolute numbers of CD4^+^ T cells (C), Foxp3^+^ Tregs (D) and neutrophils (E) in the spleen, mesenteric LNs, small intestine (SI) and colon of *Rag1*^-/-^ recipient mice 7 to 9 weeks after transfer of WT or Tafazzin-deficient T cells; means ± SEM of 3-4 host mice. Statistical analyses in (A-E) by unpaired Student’s t-tests. *, p<0.05; **, p<0.01, ***, p<0.001.

**Figure S5.**
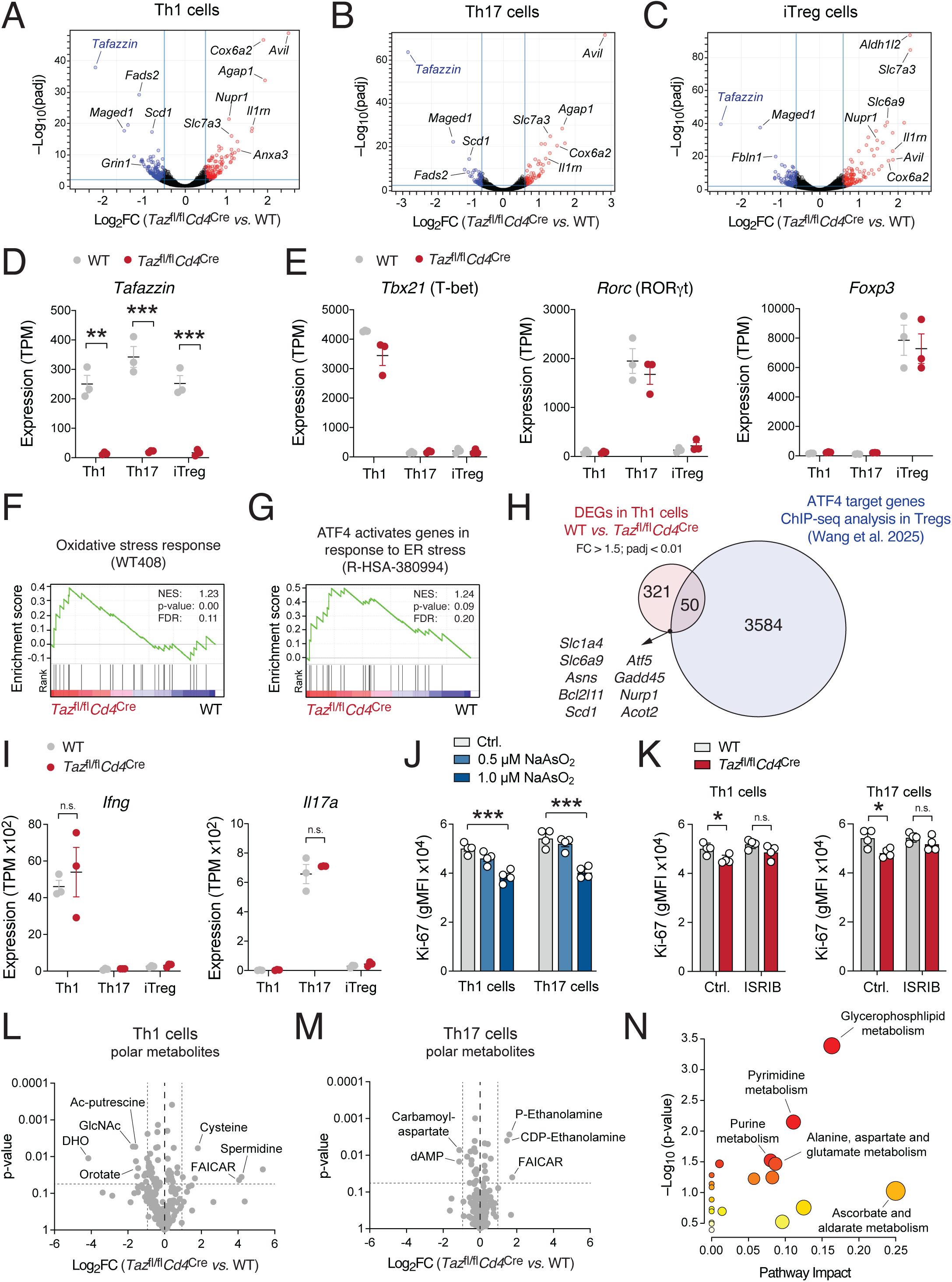
Transcriptomic and metabolomic characterization of Tafazzin-deficient T cells. (A-G) Transcriptional profiling of WT and Tafazzin-deficient T cells by RNA-sequencing (RNA-seq). **(A-C)** Volcano plots of differentially expressed genes (DEG) in WT *versus* Tafazzin-deficient Th1 (A), Th17 (B) and iTreg cells (C); 3 biological replicates per group. Genes significantly (padj < 0.01) up-and downregulated are depicted in red and blue, respectively. **(D** and **E)** Analysis of *Tafazzin* (D), *Tbx21*, *Rorc*, *Foxp3* (E) gene expression in WT and Tafazzin-deficient Th1, Th17 and iTreg cells by RNA-seq; means ± SEM of 3 biological replicates per group. **(F** and **G)** Gene set enrichment analyses (GSEA) of *oxidative stress response* (F) and *ATF4 activates genes in response to ER stress* (G) gene signatures in WT *versus* Tafazzin-deficient Th1 cells. **(H)** Overlap of DEGs between WT and Tafazzin-deficient Th1 cells with ATF4 target genes identified by ChIP-seq analyses in Treg cells (Wang et al., 2025). **(I)** Analysis of *Ifng* and *Il17a* gene expression in WT and Tafazzin-deficient Th1, Th17 and iTreg cells by RNA-seq; means ± SEM of 3 biological replicates per group. **(I)** Flow cytometric analysis of Ki-67 expression in Th1 and Th17 cells treated with 0.5 and 1 µM NaAsO_2_ for 48h; means ± SEM of 4 mice. **(J)** Analysis of Ki-67 expression in WT and Tafazzin-deficient Th1 and Th17 treated with or without 500 nM ISRIB; means ± SEM of 4 mice. **(L-M)** Metabolomic analysis of WT and Tafazzin-deficient T cells using liquid chromatography and mass spectrometry (LC/MS). **(I** and **J)** Volcano plots of differential metabolites levels between WT *versus* Tafazzin-deficient Th1 (I) and Th17 cells (J) analyzed by LC/MS; 3 biological samples per group. **(K)** Metabolic pathway enrichment analysis of differential metabolites (p < 0.05) between WT and Tafazzin-deficient T cells using MetaboAnalyst. Statistical analyses in (D), (J) and (K) by unpaired Student’s t-tests. *, p<0.05; **, p<0.01; ***, p<0.001.

**Figure S6.**
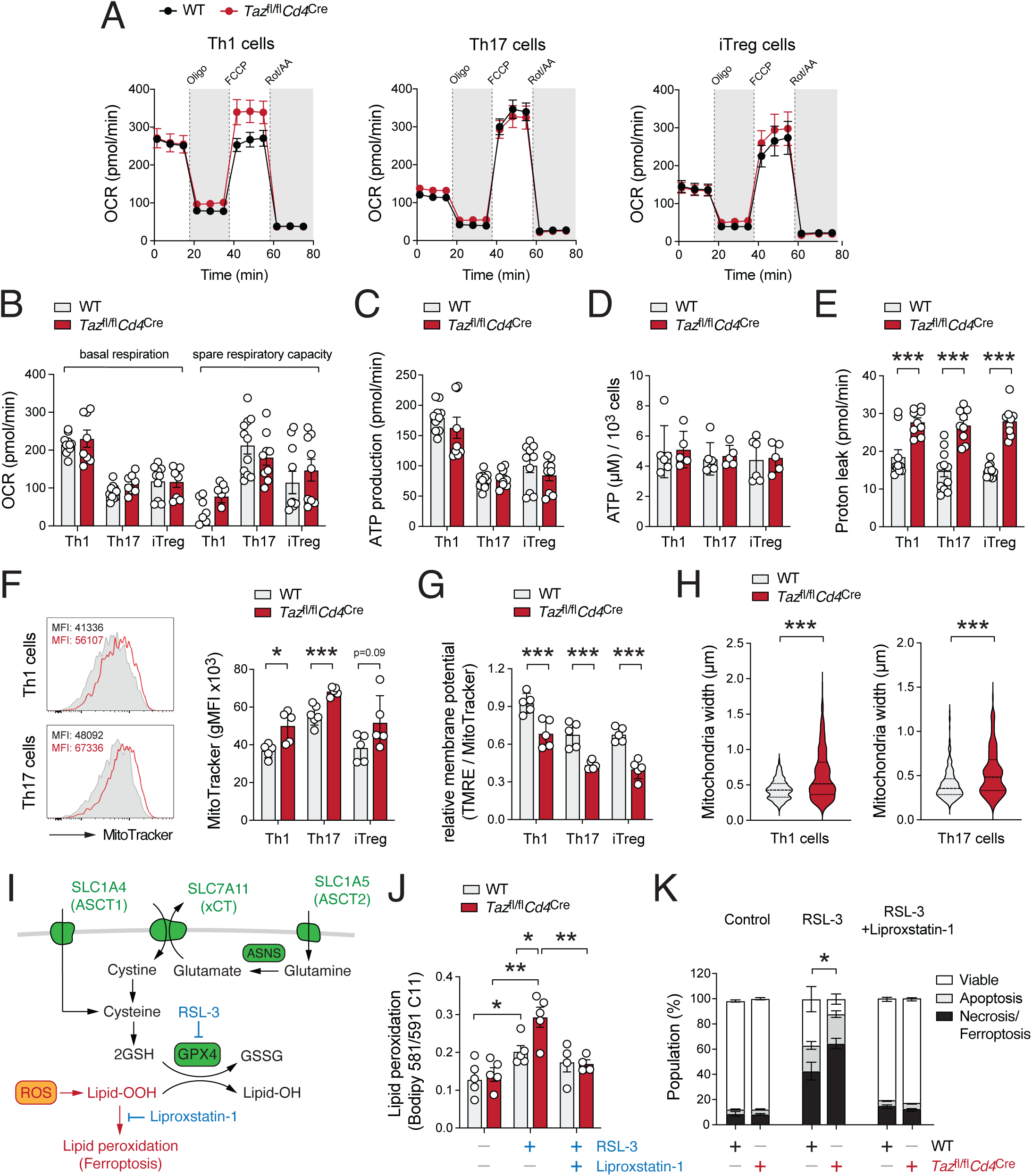
Mitochondrial performance in Tafazzin-deficient CD4^+^ T cells. **(A)** Analysis of oxygen consumption rate (OCR) in WT and Tafazzin-deficient Th1, Th17 and iTreg cells using a Seahorse extracellular flux analyzer; means ± SEM of 9-11 samples. **(B)** Quantification of basal respiration and spare respiratory capacity (SRC) in WT and Tafazzin-deficient Th1, Th17 and iTreg cells as shown in (A); means ± SEM of 9-11 samples. **(C)** ATP production rate (pmol/min) in WT and Tafazzin-deficient Th1, Th17 and iTreg cells calculated from Seahorse extracellular flux analyses; means ± SEM of 9-11 samples. **(D)** Analysis of cellular ATP concentration per 1×10^3^ cells in WT and Tafazzin-deficient Th1, Th17 and iTreg cells; means ± SEM of 5-6 samples. **(E)** Proton leak in WT and Tafazzin-deficient Th1, Th17 and iTreg cells calculated by (oxygen consumption rate after oligomycin treatment) – (non-mitochondrial respiration) of extracellular flux analyses; means ± SEM of 9-11 samples. **(F)** Analysis of mitochondrial content/mass in WT and Tafazzin-deficient Th1, Th17 and iTreg cells by flow cytometry using MitoTracker; means ± SEM of 5-6 samples. **(G)** Analysis of the relative mitochondrial membrane potential (TMRM / MitoTracker) in WT and Tafazzin-deficient Th1, Th17 and iTreg cells; means ± SEM of 5 samples. **(H)** Quantification of mitochondrial size (width) by transmission electron microscopy (TEM) as shown in Fig. 6D; analysis of > 25 individual Th1 and Th17 cells from two independent experiments. **(I)** Metabolic stress response mechanism in Tafazzin-deficient T cells. The system xCT-antiporter imports cystine in exchange for glutamate, while SLC1A4 and SLC1A5 facilitate the uptake of cysteine and glutamine, respectively, to support the biosynthesis of glutathione (GSH). GSH serves as a critical cofactor for GPX4, which converts toxic phospholipid hydroperoxides (L-OOH) into benign lipid alcohols (L-OH). The small molecule RSL-3 promotes ferroptosis by inhibiting GPX4, whereas Liproxstatin-1 counteracts ferroptosis by preventing lipid peroxidation. Transporters and enzymes highlighted in green were significantly upregulated in Tafazzin-deficient T cells (as shown in Fig. 5). **(J)** Flow cytometric analysis of lipid peroxidation in WT and Tafazzin-deficient Th1 and Th17 cells after treatment with the GPX4 inhibitor RSL-3 using Bodipy 581/591 C11 by flow ctometry; means ± SEM of 4-5 mice. **(K)** Quantification of apoptotic and necroptotic/ferroptotic cell death in WT and Tafazzin-deficient Th17 cells after treatment with the GPX4 inhibitor RSL-3 using flow cytometry as shown in Fig. 6H. Statistical analyses in (B-K) by unpaired Student’s t-tests. *, p<0.05; **, p<0.01, ***, p<0.001

**Figure S7.**
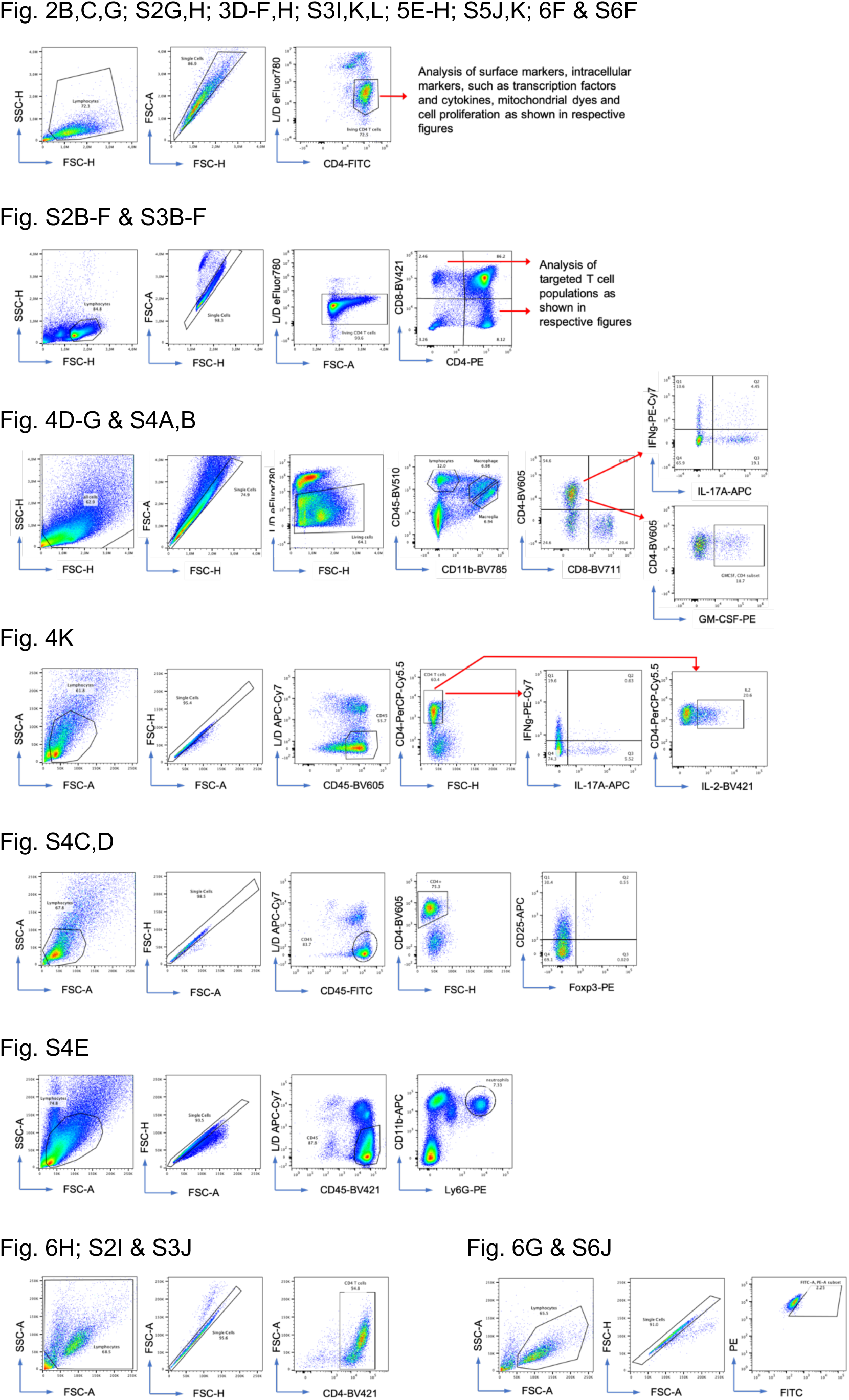
Flow cytometric gating strategies used in this study. Gating strategies for all flow cytometric analyses presented in the main and supplementary figures of this manuscript.

---

**Table S1.** List of antibodies used in this study.

| Antibody | Conjugate | Clone | Source | Cat# | RRID |
| --- | --- | --- | --- | --- | --- |
| Anti-Tafazzin | unconjugated | 2G3F7 | Lu *et al.* 2004 <sup>73</sup> | N/A | N/A |
| Anti-ATP5A | unconjugated | 15H4C4 | Thermo Scientific | 45-8099 | AB_2533835 |
| Anti-UQCRC2 | unconjugated | 13G12AF12BB11 | Thermo Scientific | 45-8099 | AB_2533835 |
| Anti-MTCO1 | unconjugated | 1D6E1A8 | Thermo Scientific | 45-8099 | AB_2533835 |
| Anti-SDHB | unconjugated | 21A11AE7 | Thermo Scientific | 45-8099 | AB_2533835 |
| Anti-NDUFB8 | unconjugated | 20E9DH10C12 | Thermo Scientific | 45-8099 | AB_2533835 |
| Anti-β-Actin | unconjugated | 2F1-1 | BioLegend | 643807 | AB_2566701 |
| Phospho-eIF2α | unconjugated | SZ01-06 | Thermo Scientific | MA5-32021 | AB_2809315 |
| Anti-Rabbit IgG | AF647 | polyclonal | Thermo Scientific | A-21244 | AB_2535812 |
| Anti-CD4 | FITC | GK1.5 | BioLegend | 100406 | AB_312691 |
| Anti-CD25 | APC | PC61 | BioLegend | 102012 | AB_312861 |
| Anti-T-bet | PE/Cyanine7 | 4B10 | BioLegend | 644824 | AB_2561761 |
| Anti-RORγt | BV421 | Q31-378 | BD Pharmingen | 562894 | AB_2687545 |
| Anti-Foxp3 | PE | FJK-16s | Thermo Scientific | 12-5773-82 | AB_465936 |
| Anti-CD4 | PerCP/Cyanine5.5 | GK1.5 | BioLegend | 100434 | AB_893324 |
| Anti-Ki-67 | PerCP/Cyanine5.5 | B56 | BD Pharmingen | 561284 | AB_10611574 |
| Anti-TNF-α | FITC | MP6-XT22 | BioLegend | 506304 | AB_315425 |
| Anti-IL-2 | BV421 | JES6-5H4 | BioLegend | 503826 | AB_2650897 |
| Anti-GM-CSF | PE | MP1-22E9 | BioLegend | 505406 | AB_315382 |
| Anti-IL-17A | APC | TC11-18H10.1 | BioLegend | 506916 | AB_536018 |
| Anti-IL-17F | AF488 | 9D3.1C8 | BioLegend | 517006 | AB_10661903 |
| Anti-IFNγ | PE/Cyanine7 | XMG1.2 | BioLegend | 505826 | AB_2295770 |
| Anti-CD4 | PE | GK1.5 | BioLegend | 100408 | AB_312693 |
| Anti-CD8a | BV421 | 53-6.7 | BioLegend | 100738 | AB_11204079 |
| Anti-CD4 | BV421 | RM4-5 | BioLegend | 100544 | AB_11219790 |
| Anti-CD45 | BV510 | 30-F11 | BioLegend | 103138 | AB_2563061 |
| Anti-CD11b | BV785 | M1/70 | BioLegend | 101243 | AB_2561373 |
| Anti-CD4 | BV605 | GK1.5 | BioLegend | 100451 | AB_2564591 |
| Anti-CD8a | BV711 | 53-6.7 | BioLegend | 100748 | AB_2562100 |
| Anti-CD45 | BV605 | 30-F11 | BioLegend | 103140 | AB_2562342 |
| Anti-CD45 | FITC | 30-F11 | BioLegend | 103108 | AB_312973 |
| Anti-CD45 | Pacific Blue | 30-F11 | BioLegend | 103126 | AB_493535 |
| Anti-CD11b | APC | M1/70 | BioLegend | 101212 | AB_312795 |
| Anti-Ly-6G | PE | 1A8 | BioLegend | 127608 | AB_1186099 |
| Anti-CD44 | FITC | IM7 | BioLegend | 103006 | AB_312957 |
| Anti-CD62L | BV510 | MEL-14 | BioLegend | 104441 | AB_2561537 |
| Anti-OX-40 | PE | OX-86 | BioLegend | 119410 | AB_2207344 |
| Anti-ICOS | AF488 | C398.4A | BioLegend | 313514 | AB_2122584 |
| Anti-PD-1 | BV605 | 29F.1A12 | BioLegend | 135220 | AB_2562616 |
| Anti-CD98 | APC | RL388 | BioLegend | 128212 | AB_2750545 |
| Anti-CD71 | PE/Cyanine7 | RI7217 | BioLegend | 113812 | AB_2203382 |
| Anti-CD69 | PE/Cyanine7 | H1.2F3 | BioLegend | 104512 | AB_493564 |

**Table S2.**
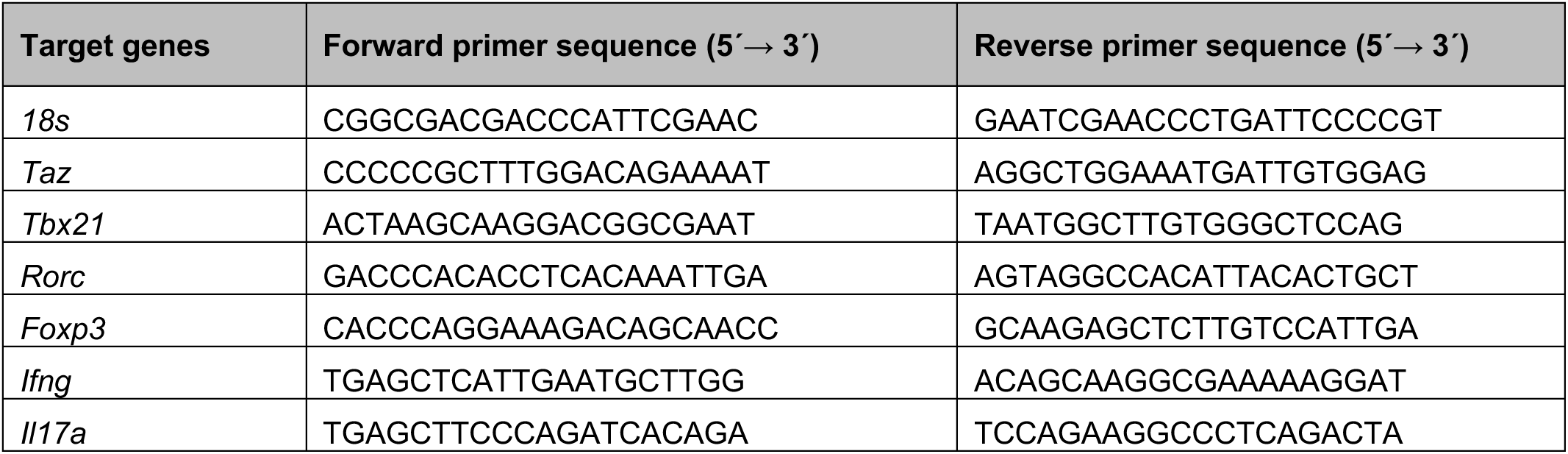
qRT-PCR primers used in this study.

